# *Panagrolaimus einhardi* sp. nov. and two sisters of fortune

**DOI:** 10.64898/2026.01.07.697871

**Authors:** Laura C. Pettrich, Arunee Suwanngam, Nadège Guiglielmoni, Noelle Ledoux, Laura I. Villegas, Amy Treonis, Lewis Stevens, Manuela Kieninger, Michael Paulini, Mark Blaxter, Ann-Marie Waldvogel, Oleksandr Holovachov, Philipp H. Schiffer

## Abstract

Identifying nematodes to the species level is known to be complicated due to their morphological plasticity and limited number of taxonomically important characters. This is especially apparent in the genus *Panagrolaimus*, which comprises many cryptic species that are morphologically difficult to distinguish but differ genetically. These roundworms are particularly notable for their adaptation to extreme environments that are inhospitable to many other forms of life. Traditional morphological identification methods often fail at distinguishing genetically divergent populations due to high morphological plasticity in *Panagrolaimus*, limiting the efficacy of species discovery. High-quality genome assemblies overcome these challenges, offering a comprehensive blueprint of an organism’s genetic structure that can be used for species identification. The analysis of ultra-conserved elements across multiple loci harvested from genome assemblies provides robust phylogenetic resolution. In this study, we integrate genome sequencing, ultra-conserved element analysis, and morphological assessment to identify and describe three novel species: *Panagrolaimus einhardi* sp. nov., formerly *Panagrolaimus* sp. ES5 from Germany; *Panagrolaimus shuimeiren* sp. nov. from the Namib Desert; and *Panagrolaimus nebliphilus* sp. nov. from the Atacama Desert. *P. einhardi* sp. nov. is named after Prof. Einhard Schierenberg, a renowned expert in roundworm development and cherished member of the nematode community, who isolated this species himself. All three species originate from different geographical locations, and their respective identification are supported by high-quality genome assemblies from either PacBio HiFi or Oxford Nanopore long-read data. The *P. einhardi* sp. nov. genome was scaffolded using Hi-C technology, which resulted in a 116 Mb collapsed assembly composed of 44 scaffolds (N50: 28 Mb). *P. shuimeiren* sp. nov. has an assembly size of 69 Mb with 49 scaffolds and a N50 of 13 Mb. *P. nebliphilus* sp. nov. assembly is 70 Mb with 24 scaffolds (N50: 13 Mb). The capacity of *Panagrolaimus* to adapt to extreme environments is driving research into their survival mechanisms, requiring comprehensive genomic resources. By combining morphology and genomics, we can gain a more comprehensive understanding of the rich biological diversity in lineages with numerous cryptic species, such as the Panagrolaimidae, thereby clarifying relationships where morphological data alone are ambiguous or confounded.

## INTRODUCTION

Nematodes are highly dominant in soil food web and contribute to a large proportion of terrestrial biomass (approximately 0.3 gigatons globally) (Bardgett and van der Putten, 2014; van den Hoogen et al., 2019), participating in carbon and nitrogen cycling (Gebremikael et al., 2016; van den Hoogen et al., 2019; Ferris et al., 1997). Various phylogenetically unrelated species of free-living nematodes were found to survive extreme conditions and inhabit environments hostile to the majority of multicellular eukaryotes and being major drivers of nutrient cycling and energy flow (McSorley, 2003). Among them is the bacteria-feeding nematode genus *Panagrolaimus*, originally described by Fuchs (1930), belonging to the suborder Tylenchina of the order Rhabditida (Clade IV as defined by Blaxter et al. (1998)). It has a cosmopolitan distribution, including a wide variety of habitats (Shannon et al., 2005), ranging from polar (Shatilovich et al., 2023; Raymond and Wharton, 2012; Boström, 1988), to temperate, subtropical and tropical to non-polar deserts (Villegas et al., 2024b; Rawson et al., 2024). They are successful colonizers, facilitated by their broad ecological tolerance, short life cycles, and high fecundity, resulting in dense populations under favorable conditions (Bongers and Ferris, 1999; Bongers, 1990). Many *Panagrolaimus* species can survive unfavorable and even extreme conditions by entering a state of suspended metabolic activity known as cryptobiosis (Cooper and Gundy, 1971). Their ability to subsist inactively over very long periods of time has been shown in *Panagrolaimus kolymaensis* Shatilovich et al., 2023 which was revived from a 46,000-year-old Russian permafrost sample (Shatilovich et al., 2023).

The statement that nematodes are morphologically uniform, just “worms”, is completely unjustified when it refers to the entire phylum, but it perfectly describes the genus *Panagrolaimus*. Few taxonomically useful morphological (qualitative) characters, high plasticity of morphometrics, ubiquitous distribution throughout nearly all terrestrial environments hides enormous uncatalogued taxonomic diversity within this genus. Morphological description remains fundamental in nematode taxonomy, but it is often insufficient in lineages with numerous cryptic and sibling species, where qualitative features overlap while quantitative diagnostic characters are subtle, and is largely inapplicable for identification of developmental stages (Mianowska, 1977; Bredtmann et al., 2017). If morphologically at all possible, nematode species description demands specialized expertise and access to comparative voucher material (Eyualem and Blaxter, 2003; Hodda, 2022). Molecular approaches, such as single- or multi-locus marker sequencing, have significantly enhanced species delineation in the last decades (Shatilovich et al., 2023; Blaxter et al., 1998). Commonly used standard markers like 18S rDNA or 28S rDNA offer limited phylogenetic resolution at species level in many recently diversified lineages, while at the same time are prone to intragenomic polymorphism (Wang et al., 2023), making their use for species delimitation and taxonomic placement problematic (Eyualem and Blaxter, 2003; Rawson et al., 2024; Villegas et al., 2025; Ahmed et al., 2022). COX1, the most commonly used barcode in many other animal lineages, is rarely used in nematode taxonomy due to it being too variable there, and often impossible to sequence even with taxon-specific primers (Scharhauser et al., 2020). As such, the rate of species description lags far behind species discovery in actively studied genera of nematodes, such as *Caenorhabditis* and *Pristionchus* (Hodda, 2022). Similarly, numerous and not-standardized strain numbers instead of formal species names for *Panagrolaimus* persist in the literature and scientific databases, highlighting the need for more detailed and accurate taxonomic classification within this genus (Hodda, 2022).

The effectiveness of marker-based species delineation in nematodes can be limited by: (i) low sequence variability, as exemplified by rRNA genes in recently diversified lineages; (ii) intragenomic polymorphism, again in rRNA genes; (iii) or high diversification rates in COX1. Thus, markers not only insufficiently capture the evolutionary history of the taxon in question (Rawson et al., 2024; Villegas et al., 2025), but also prevent the implementation of universal phylum-wide criteria for species delimitation. Whole-genome sequencing offers a more comprehensive and unbiased approach that encompasses both coding and non-coding regions, structural variants, and patterns of conservation and divergence across the genome (Theissinger et al., 2023; Ahmed et al., 2022). The application of genomic data has significantly refined phylogenetic analyses within the genus *Caenorhabditis*, particularly given the high morphological conservation. An early example of using genomic information to clarify relationships in *Caenorhabditis* is Slos et al. (2017), which examined *Caenorhabditis monodelphis* and combined morphology with a new draft genome to determine its position within the genus. By analyzing hundreds of single-copy orthologs from *C. monodelphis*, they reconstructed a molecular phylogeny that positioned the species as a basal lineage, making its genome vital for species identification and understanding deep evolutionary relationships. Stevens et al. (2019) expanded the genomic approach by providing draft genomes for nine new *Caenorhabditis* species and one transcriptome assembly and integrating them into a phylogenomic analysis where they identified nearly 2,000 single-copy orthologs across 32 species and reconstructed a highly resolved phylogeny. Unlike earlier multi-gene studies, this genome-wide approach offers greater resolution. The study also found that genome size and developmental genes relate to phylogenetic position, making genomes valuable for understanding evolutionary relationships.

One approach for species delineation is to extract ultra-conserved elements (UCEs) (Bejerano et al., 2004) and the specific variability of their flanking regions from genome assemblies to increase taxonomic resolution and delineate species (Faircloth et al., 2012). Commonly used in other invertebrate groups (Faircloth et al., 2014; Starrett et al., 2016; Blaimer et al., 2015; González-Delgado et al., 2024), this approach was also successfully applied for *Panagrolaimus* nematodes (Villegas et al., 2025). There is a high potential for the use of high-quality genomes as a transformative tool for species delineation and identification (Ahmed et al., 2022) and to tackle complex taxonomic and phylogenetic questions in the diverse but understudied phylum Nematoda. However, the availability of high-quality genome assemblies with correctly identified and accessible reference species is fundamental.

As *Panagrolaimus* is emerging as a (model) system to study the evolutionary genomics of life traits and ecological function in soil ecosystems, there is a necessity to refine taxonomic classification in the genus (Schiffer et al., 2019; Lewis et al., 2009; Villegas et al., 2024a, 2025; Shatilovich et al., 2023; Tyson et al., 2012). *Panagrolaimus* has been utilized as a model for investigating the evolution of reproductive modes. Two different reproduction modes can be found in the genus *Panagrolaimus*: gonochoristic (female/male), and parthenogenetic (Lewis et al., 2009). Parthenogenesis in *Panagrolaimus* is suspected to have originated from a hybridization event 1.3-8.5 million years ago, which changed the ploidy from diploid to triploid (Schiffer et al., 2019). A potential separate hybridization event may have occurred for *P. kolymaensis* (Shatilovich et al., 2023). Furthermore, the occurrence of *Panagrolaimus* in extreme and disturbed environments, along with their variable reproductive modes, makes them especially valuable for studying genome evolutionary processes and understanding the limits of life.

*P. einhardi* sp. nov., previously referred to as *Panagrolaimus* sp. strain ES5 and *Panagrolaimus* sp. bornheim, has been established as a long-term laboratory model species and was first mentioned by Lewis et al. (2009). Studies on parthenogenesis and cryptobiosis (Schiffer et al., 2019), as well as mutations (Villegas et al., 2024a), have been published using ES5 as a test subject, but it lacked a proper species name and taxonomic description until now. The other two species, *P. shuimeiren* sp. nov. and *P. nebliphilus* sp. nov., originate from two westward directed coastal desert systems: the Namib Desert of Namibia and the Atacama Desert of Chile respectively. The Atacama Desert is described as the oldest non-polar desert on Earth and was formerly depicted as devoid of life, but has revealed a rich biodiversity in spite of its extreme conditions (Gómez-Silva and Batista-García, 2022; Hartley et al., 2005), which is also found for nematodes (Villegas et al., 2024b). The Namib Desert hosts a variety of nematode species, though most studies have historically focused on Cephalobidae (Rashid et al., 1990a; Rashid and Heyns, 1990b; Rashid et al., 1990b; Rashid and Heyns, 1990a), including several *Chiloplacus* species (Rashid and Heyns, 1990a) and the genus *Namibinema* (Rashid and Heyns, 1990b). Other families, like Belonolaimidae, have also been recorded (Rashid and Heyns, 1990c). More recently, attention has turned to Panagrolaimidae, with the discovery of a new species, *Panagrolaimus namibiensis* (Rawson et al., 2024). Environmental factors, such as fog and sporadic rainfall, appear to influence nematode activity: the study observed that nematodes emerge from anhydrobiosis and become active during fog events, and that diversity tends to increase along gradients of decreasing fog (Treonis et al., 2022). Additionally, the endemic plant *Welwitschia mirabilis* has been found to support diverse nematode communities in the Namib Desert, with *Panagrolaimus* as the most abundant species documented (Treonis et al., 2024).

Here, we set a new standard by describing three new gonochoristic species of *Panagrolaimus* using an integrative approach that combines detailed morphological examination using light and scanning electron microscopy supported by the genome assemblies, utilizing PacBio HiFi coupled with Hi-C or Oxford Nanopore sequencing, followed by the phylogenetic analysis based on ultra-conserved elements, in order to resolve species boundaries of newly described taxa and their interrelationships with other congeners within *Panagrolaimus*, demonstrating the value of genomic resources for accelerating and strengthening species descriptions in nematodes.

## MATERIAL AND METHODS

### Sample collection and culturing

*P. einhardi* sp. nov. was collected from the soil around a dead blackberry (*Rubus* spp.) in Bornheim, Germany by Prof. Einhard Schierenberg in 2006 (Schiffer et al., 2014) and has been maintained in the laboratory culture for many years. The other two species have been retrieved from soil samples from two desert systems. *Panagrolaimus shuimeiren* sp. nov. (WWL115) originates from a sample campaign to the Namib Desert. Fieldwork and sampling were conducted in compliance with all necessary permits and permissions granted by the National Commission on Research, Science and Technology (Permit Number: RPIV01042034). *Panagrolaimus nebliphilus* sp. nov. (WWL072) originates from a sampling campaign to the Atacama Desert and did not require any sampling permits.

Nematodes were extracted from soil using seedling trays (sprouting) as described in Villegas et al. (2024b), which is a modification of a widely known Whitehead and Hemming tray extraction method (Whitehead and Hemming, 1965). Specifically, seeding trays were lined with a single layer of Kimberly-Clark Kimtech Science Precision Wipes 7551 (Kimberly-Clark, Irving, TX, USA) in the upper compartment, and the soil sample was dispersed thinly (no more than 1 cm) on top of the paper. The lower compartment of the trays was filled with water, and the samples were left to rehydrate overnight, and allow nematodes to move downwards. The water from the lower compartment was subsequently filtered through a 45, 80 or 120 µm sieves to concentrate nematodes, which were transferred into a counting or petri dishes and analyzed under a Zeiss Stemi 2000 microscope (Zeiss, Oberkochen, Baden-Württemberg, Germany). Some of the extracted nematodes were picked out from the suspension and placed on 2% plain agar plates with 0.1 % cholesterol and inoculated with *Escherichia coli* OP50, and maintained continuously under controlled conditions.

### Morphological characterization

Nematodes were studied both live and preserved on permanent slides. For live observations, nematodes were mounted on temporary slides using food-grade silicon as a support for the cover slip (Kanzaki, 2013), and subsequently observed/photographed using Nikon Eclipse 80i microscope equipped with differential interference contrast (DIC) optic and digital Sony a6400 camera. Photographs were processed using Capture One 12 and Adobe Photoshop Elements. For permanent mount, live nematodes were first collected in distilled or deionized water, concentrated and fixed in an equal volume of hot (60-80 °C) fixative solution following the Seinhorst method (Seinhorst, 1959). Specimens were kept for 1-2 days at 4 °C to ensure thorough fixation. Fixed nematodes were processed into pure glycerin by slowly evaporating the ethanol and increasing the glycerin proportion using the glycerin-ethanol method. Individual nematodes were mounted in glycerin drops and sealed with paraffin wax on permanent slides. Mounted nematodes were examined using various microscopes equipped with DIC optics. Imaging and morphometric measurements were conducted using the stream essentials system and the AxioVision software. Measurements were recorded, and the average and standard error were calculated.

### Scanning electron microscopy

Live nematodes were washed from the agar plates and fixed in a mixture of 4 % paraformaldehyde and 2 % glutaraldehyde in 0.1 M sodium cacodylate buffer, followed by a secondary fixation in 1 % osmium tetroxide, and subsequently run through serial ethanol dehydration. The final suspension of nematodes in 100 % ethanol was critical-point dried, mounted, and sputter-coated with 10 nm gold palladium. Images were taken using a JEOL JSM-IT700HR scanning electron microscope (JEOL USA, Inc., Peabody, MA, USA).

### Ultra-low input PacBio HiFi sequencing

DNA for ultra-low input PacBio HiFi sequencing was prepared as described in Guiglielmoni et al. (2024). Briefly, up to 10 *Panagrolaimus einhardi* sp. nov. individuals were collected from single female progeny and washed in water, flash-frozen using liquid nitrogen in a salt-based extraction buffer (Tris-HCl 100 mM, ethylenediaminetetraacetic acid 50 mM, NaCl 0.5 M and sodium dodecylsulfate 1%) and incubated overnight at 50°C with 5 µL of proteinase K (Product code D3001-2, Zymo Research, Irvine, CA, USA). DNA was precipitated using NaCl 5 M, yeast tRNA and isopropanol, and washed twice with 80% ethanol before elution in a commercial elution buffer (Product code D3004-4-10, Zymo Research, Irvine, CA, USA). RNA was removed using RNAse (Product code 19101, Qiagen, Hilden, North Rhine–Westphalia, Germany) for 1 hour at (37°C). DNA concentrations were quantified using a Qubit 4 fluoremeter (Thermo Fisher Scientific, Waltham, MA, USA) with 1X dsDNA kit. HiFi libraries were prepared with the Express 2.0 Template kit (Pacific Biosciences, Menlo Park, CA, USA) and sequenced on a Sequel II/Sequel IIe instrument with 30 hours movie time. PacBio HiFi reads were generated using SMRT Link (v10, Pacific Biosciences, Menlo Park, CA, USA) with default parameters.

### Oxford Nanopore sequencing

Nematodes of *P. shuimeiren* sp. nov. and *P. nebliphilus* sp. nov. were each washed using autoclaved and deionized water from agar plates and pelleted at 3,000 g for 5 min. Samples were decontaminated using a sucrose flotation method by resuspending the worms with a 1 M sucrose solution and centrifugation at 4,500 x g for 5 min, and transfer of 2-3 ml of the upper layer. The solution was diluted with nuclease-free water, and worms were pelleted again at 3,000 x g for 5 min. Pellets were immediately used or frozen at −20 °C until further use. Genomic DNA was extracted from pellets using the Monarch HMW DNA Extraction Kit for Tissue from NEB (Product code T3060L, New England Biolabs, Ipswich, MA, USA) as described in Kieninger (2025). Initially, 600 µL of HMW gDNA Tissue Lysis Buffer was mixed with 20 µL of Proteinase K to create the Lysis Mix. A portion of this buffer (50 µL) was added to a frozen nematode pellet on ice, followed by pestle homogenization with at least 30 up-and-down movements in one direction. After adding the remaining Lysis Mix, the sample was digested at 56 °C in the ThermoMixer at 750 rpm for 15 min, then at 550 rpm for 2 hours. RNA digestion was done by adding 10 µL of RNase A and incubating at 56 °C, 550 rpm for 10 min. The samples were centrifuged at 2,500 x g for 5 min, and the supernatant was collected. Following that, 300 µL of cold Monarch Protein Separation Solution was added, and the samples were inverted for 3 min and incubated on ice for 10 min before being centrifuged at 16,000 x g for 25 min. The upper phase was collected, and three Monarch DNA Capture Beads were added. After mixing with 550 µL of isopropanol and washing with Monarch gDNA Wash Buffer, the glass beads were transferred to a new tube. Elution was performed in two parts with 100 µL and 150 µL of Monarch gDNA Elution Buffer II at 56 °C for 10 min each, followed by a centrifuge spin at 12,000 x g for the second elution and further incubation at 37 °C with shaking at 300 rpm for 30 min. Libraries for Oxford Nanopore sequencing were prepared using the Ligation Sequencing Kit LSK114 (Product code SQK-LSK114, Oxford Nanopore Technologies, Oxford, UK). Sequencing was performed on a R10.4.1 PromethION flowcell (Product code FLO-PRO114M, Oxford Nanopore Technologies, Oxford, UK). Reads were basecalled using Dorado vX.X.X (Oxford Nanopore Technologies, 2022) in duplex mode with model dna r10.4.1 e8.2 400bps supvX.X.X and the reads were converted to fastq using SAMtools v1.6 (Danecek et al., 2021) with the module samtools fastq.

### Hi-C sequencing

Pellets of frozen *Panagrolaimus einhardi* sp. nov. nematodes weighing approximately 20 mg were processed using the Arima Hi-C version 2 kit (Arima Genomics, Carlsbad, California, USA) following the manufacturer’s instructions. Illumina libraries were prepared using the NEBNext Ultra II DNA Library Prep Kit (Product code E7645L, New England Biolabs, Ipswich, MA, USA) and each library was sequenced on one-eighth of a NovaSeq S4 lane using paired-end 150 bp sequencing. Hi-C library preparation and sequencing were performed by the Scientific Operations: Sequencing Operations core at the Wellcome Sanger Institute.

### RNA sequencing

RNA was extracted from pellets of frozen *Panagrolaimus einhardi* sp. nov. nematodes weighing at least 20 mg. We suspended the frozen nematodes in 750 µL of TRIzol (Invitrogen, Thermo Fisher Scientific, Waltham, MA, USA), which was flash frozen in liquid nitrogen before being thawed completely. This freezing/thawing process was repeated a total of three times. The RNA was extracted following the manufacturer’s protocol. RNA was eluted in 100 µL MagMAX Total RNA Elution Buffer (Thermo Fisher Scientific, Waltham, MA, USA). The RNA was digested with DNase using the TURBO DNA-freeTM Kit (Invitrogen, Thermo Fisher Scientific, Waltham, MA, USA). Quality control was performed using the NanoDrop One photospectrometer (Thermo Fisher Scientific, Waltham, MA, USA) and the Tapestation 4150 (Agilent Technologies, Santa Clara, CA, USA) with the RNA tape. The RNA had RIN values of between 7.2 and 10. Concentrations were confirmed by Qubit measurements (Thermo Fisher Scientific, Waltham, MA, USA). Illumina libraries were prepared using the Stranded RNAseq Kit with an Oligo dT pulldown and sequenced in 96-plex on a NovaSeq S4 lane using paired-end 150 bp sequencing. RNA library preparation and sequencing were performed by the Scientific Operations: Sequencing Operations core at the Wellcome Sanger Institute.

### Nuclear genome assemblies

PacBio HiFi reads were assembled using hifiasm v0.25.0 (Cheng et al., 2021) with parameter −l 3. For Nanopore reads, the option `--ont` was used. Long reads were mapped against the draft assemblies using minimap2 v2.24 (Li, 2018) with parameters `-ax map-hifi` for PacBio HiFi reads and `-ax map-ont` for Nanopore reads, then the output was sorted using SAMtools v1.6 with the module samtools sort. Contigs were aligned against the nt database using BLAST v2.13.0 (Altschul et al., 1990). Orthologs were searched for using the Benchmarking Universal Single-Copy Orthologs (BUSCO) (Manni et al., 2021b,a) tool v5.4.7 against the Metazoa odb10 lineages with parameter `-m genome`. Mapped reads, BLAST hits and BUSCO orthologs were provided as input to BlobToolKit v4.1.5 (Challis et al., 2020), and contaminants were subsequently removed. Long reads were mapped again against the decontamined assemblies using minimap2 v2.24 with parameter `-x map-hifi` or `-x map-ont`. The tool purge dups v1.2.5 (Guan et al., 2020) was used to remove uncollapsed haplotypes. Hi-C reads were were trimmed using Trim Galore v0.6.10 (Krueger et al., 2023), mapped using hicstuff v3.1.5 (Matthey-Doret et al., 2020) and bowtie2 v2.2.5 (Langmead and Salzberg, 2012) with parameters `-e DpnII,HinfI -m iterative`, and the processed output was used for scaffolding with instaGRAAL v0.1.6 no-opengl (Baudry et al., 2020) with parameters −l 4 -n 100 -c 0 `-N` 5. The chromosome-level assembly of *Panagrolaimus einhardi* sp. nov was checked for residual contamination and corrected using BlobToolKit (Challis et al., 2020) and FCS (Astashyn et al., 2024). Manual curation was performed using JBrowse 2 (Diesh et al., 2023), HiGlass (Kerpedjiev et al., 2018) and PretextView (Harry, 2022). K-mer analyses were conducted using Kmer Analysis Toolkit (KAT) v2.4.2 (`kat comp`) with k-mer sizes specified (`k` = 27). Sequencing reads (PacBio Hifi or Oxford Nanopore) were compared against the assembled genome. To estimate the completeness of the assemblies, BUSCO v5.8.2 was utilized using the Nematoda odb12 lineage (Tegenfeldt et al., 2024; Manni et al., 2021b,a). The summary of the assemblies were visualized in snail and blob plots using BlobToolKit (Challis et al., 2020). For the blob plots the reads were re-mapped against the genome assemblies using minimap2 v2.28 and to identify potential taxonomic affiliations of assembled scaffolds, sequence similarity searches were run against the NCBI nucleotide database using BLASTN. The genome assembly was queried against a locally installed copy of the NCBI nt database using BLAST+ v2.17.0 (`-outfmt “6 qseqid staxids bitscore std” -max target seqs 10 -max hsps 1 -evalue 1e-25`). The three assemblies were compared and visualized using Circos v0.69-9 (Krzywinski et al., 2009). The assemblies were pairwise aligned using minimap2 v2.28 (Li, 2018), accommodating up to approximately 10 % sequence divergence (`-x asm10`), resulting in PAF alignment files. Scaffold coordinates were extracted to BED files. Based on this a Circos karyotype file was created using the mapping, with unique colors designated for each assembly. The pairwise alignments were filtered to keep only the main scaffolds. Distinct link files were created for each assembly pair.

### Mitochondrial genome assemblies

The module `findMitoReference.py` from MitoHiFi v3.2.1 (Uliano-Silva et al., 2023) was run with parameter `--type mitochondrion` to search for a closely related mitochondrial genomes to use as a reference and identified the complete mitochondrial genome of *Panagrellus redivivus* (AP017464) (Kikuchi et al., 2019). MitoHiFi was run with reads as input (`-r`) and parameters `-o 5 --mitos`.

### Repeat and gene annotation

The Extensive *De novo* TE Annotator (EDTA) pipeline v2.0.1 (Ou et al., 2019) was run with parameters `--sensitive 1 --anno 1 --force 1`. The hardmasked assembly was converted into a softmasked assembly. For *Panagrolaimus einhardi*, RNA reads were trimmed using TrimGalore v0.6.10 (Krueger et al., 2023) and mapped to the assemblies using hisat2 v2.2.1 (Kim et al., 2019). After sorting the mapped reads using SAMTools v1.6 (Danecek et al., 2021), the output was provided as input to BRAKER v3.0.3 (Gabriel et al., 2023) with parameters `--gff3 --UTR off`, which combined predictions from Augustus v3.5.0 (Stanke et al., 2008) and GeneMark-ES v4.68 (Lomsadze et al., 2005). For species for which no RNA-seq data were available, predicted genes were provided as protein hints. The longest isoform was selected using Another Gtf/Gff Analysis Toolkit (AGAT) v0.8.0 (Dainat, 2020) with the script `agat_sp_keep_longest_isoform.pl`.

### Phylogenetic reconstruction using ultra-conserved elements (UCEs)

Phylogenetic reconstruction was performed using the master probe list and following the methods of Villegas et al. (2025), and this tree was also used as a reference. Genomes of the Panagrolaimidae family accessible via ENA were used for a phylogenetic reconstruction using ultra-conserved Elements (UCEs) (Supplementary Material Table S1). The software PHYLUCE v1.7.3 (Faircloth, 2015; Faircloth et al., 2012) was employed according to Tutorial III and then Tutorial I. The genome assemblies were transformed into 2bit format using faToTwoBit v482 (Kuhn et al., 2012). UCE probes from Villegas et al. (2025) were aligned with the genomes using `phyluce_probe_run_multiple_lastzs_sqlite (--identity 50)`. The corresponding FASTA sequences were extracted using `phyluce_probe_slice_sequence_from_genomes (--flank 500)` to harvest the UCEs from the genomes. UCE loci obtained from genomes were matched to probes with `phyluce_assembly_match_contigs_to_probes`. These were then extracted with `phyluce_assembly_get_match_counts` and phyluce assembly get fastas from match counts. UCEs for the species were aligned with `phyluce_align_seqcap_align` and afterwards trimmed with `phyluce_align_get_trimal_trimmed_alignments_from_untrimmed`. A summary of the alignment was calculated with `phyluce_align_get_align_summary_data`. Afterwards, the alignment was cleaned with `phyluce_align_remove_locus_name_from_files` and to create a data matrix the command `phyluce_align_get_only_loci_with_min_taxa (--percent 0.75)` was utilizied. The final alignment was concatenated (phyluce align concatenate alignments). With the final alignment, a tree was created using IQ-TREE v2.2.2.7 (Minh et al., 2020) with the settings to automatically find the best-fit substitution model (`-m MFP`) and perform 1000 ultrafast bootstrap replicates (-bb 1000). The tree was visualized and finalized in TreeViewer v2.2.0 (Bianchini and Sánchez-Baracaldo, 2024), where the tree was rooted, taxa were renamed and colored, and bootstrap values were added. Four *Acrobeloides* species were used as outgroup: *A. obliquus*, *A. thornei*, *A. tricornis* PAP.22.17 (WWL709) and *A. maximus*.

### Phylogenetic reconstruction using 18S rDNA

A phylogenetic tree was reconstructed based on Abolafia (2025), and corresponding reference sequences were downloaded from NCBI Gen Bank. Additionally, 18S rDNA sequences were added from the genome assemblies, which were also to build the UCE phylogeny. To obtain the sequences, first the 2bit format was converted back to FASTA (`twoBitToFa`) and the tool barrnap v0.9 (Seemann, 2018) was employed specifying the kingdom eukaryotes (`--kingdom euk`). Matching regions were extracted as FASTA file. Extracted sequences with a minimum length of 300 bp were manually checked, and if there were several sequences for the same assembly, a consensus sequence was created by aligning them using MAFFT v7.110 (`--auto`) (Katoh and Standley, 2013) and a consensus was created using majority rule approach. For *P. davidi* no sequence was found using barnap, so instead a 18S sequence from NCBI of *P. davidi* (accession: AJ567385) was blasted locally against the assembly (`makeblastdb -dbtype` nucl and `blastn -outfmt “6 qseqid sseqid pident length mismatch gapopen qstart qend sstart send value bitscore sseq” -max_target_seqs 1 -evalue 1e-20`) to extract the corresponding 18S sequence. Sequences with a minimum length of 600 bp were used to create a multi-species alignment using the MUSCLE algorithm as implemented in AliView (Larsson, 2014). The resulting alignment was used to reconstruct a phylogenetic tree in IQTree v3.0.1 (`-st DNA -m MFP -msub nuclear -b 1000 -T AUTO -o Acrobeloides_maximus`) (Minh et al., 2020). The tree was visualized and finalized in TreeViewer v2.2.0 (Bianchini and Sánchez-Baracaldo, 2024), where the tree was rooted, names of taxa were edited and colored, and bootstrap values were added. The same species were used as outgroup as in the UCE-based phylogeny.

### Comparison of orthologous clusters

Protein sequences of the assemblies were clustered in OrthoVenn3 (Sun et al., 2023) using the OrthoFinder algorithm and standard settings with an e-value of 1e-2 and an inflation value of 1.50. Colors were adapted using Inkscape v.1.3.2.

## RESULTS

### Descriptions

> Phylum Nematoda Potts, 1932
>
> Class Chromadorea Inglis, 1983
>
> Suborder Tylenchina Thorne, 1949
>
> Family Panagrolaimidae Thorne, 1937
>
> Genus Panagrolaimus Fuchs, 1930
>
> Panagrolaimus einhardi sp. nov.
>
> (Figures 1A-B, 2-3; Table 1)

**Figure 1.**
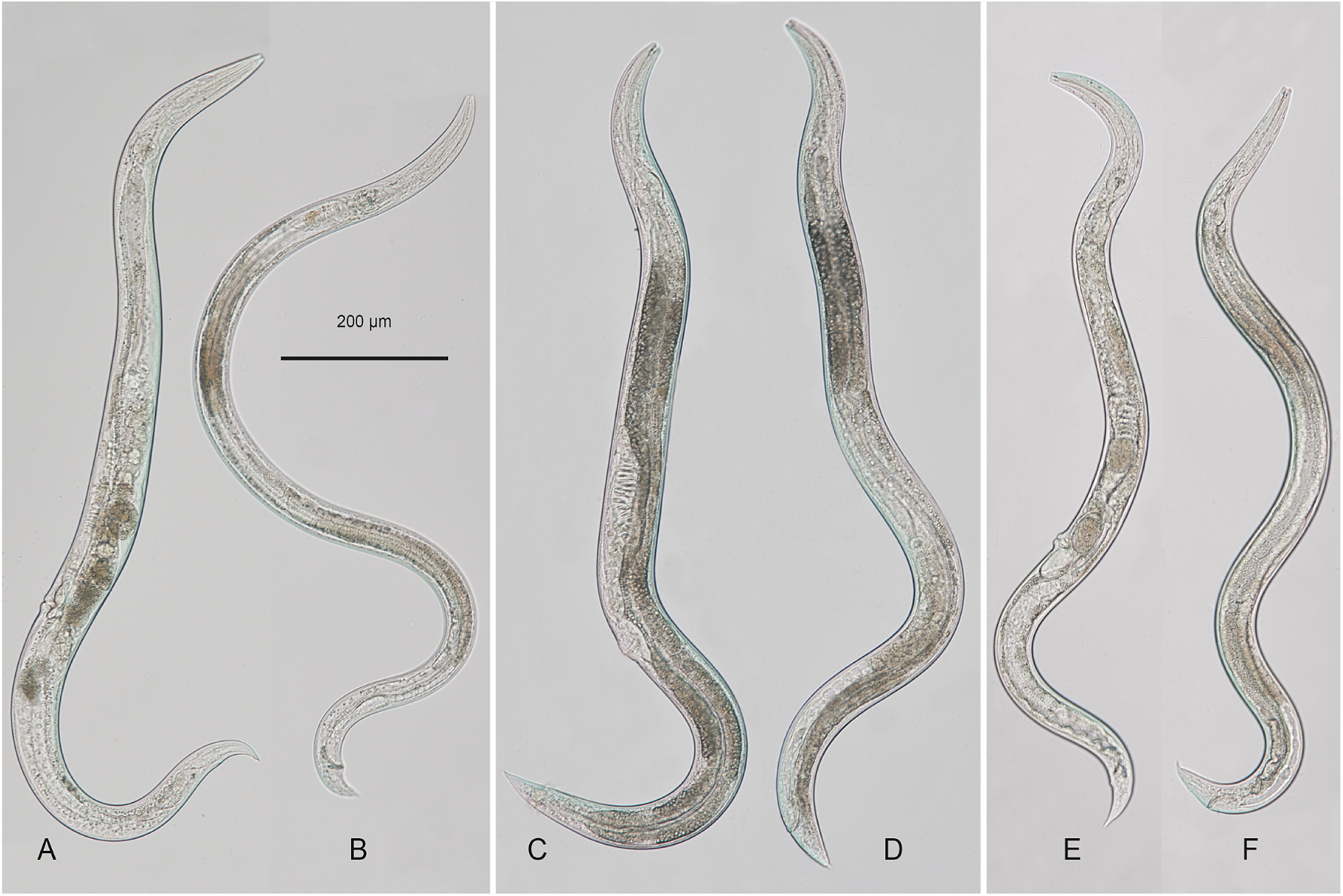
Whole-body light microscopy images of ***Panagrolaimus einhardi*** sp. nov. (A, female. B, male), ***Panagrolaimus shuimeiren*** sp. nov. (C, female, D, male) and ***Panagrolaimus nebliphilus*** sp. nov. (E, female. F, male).

**Figure 2.**
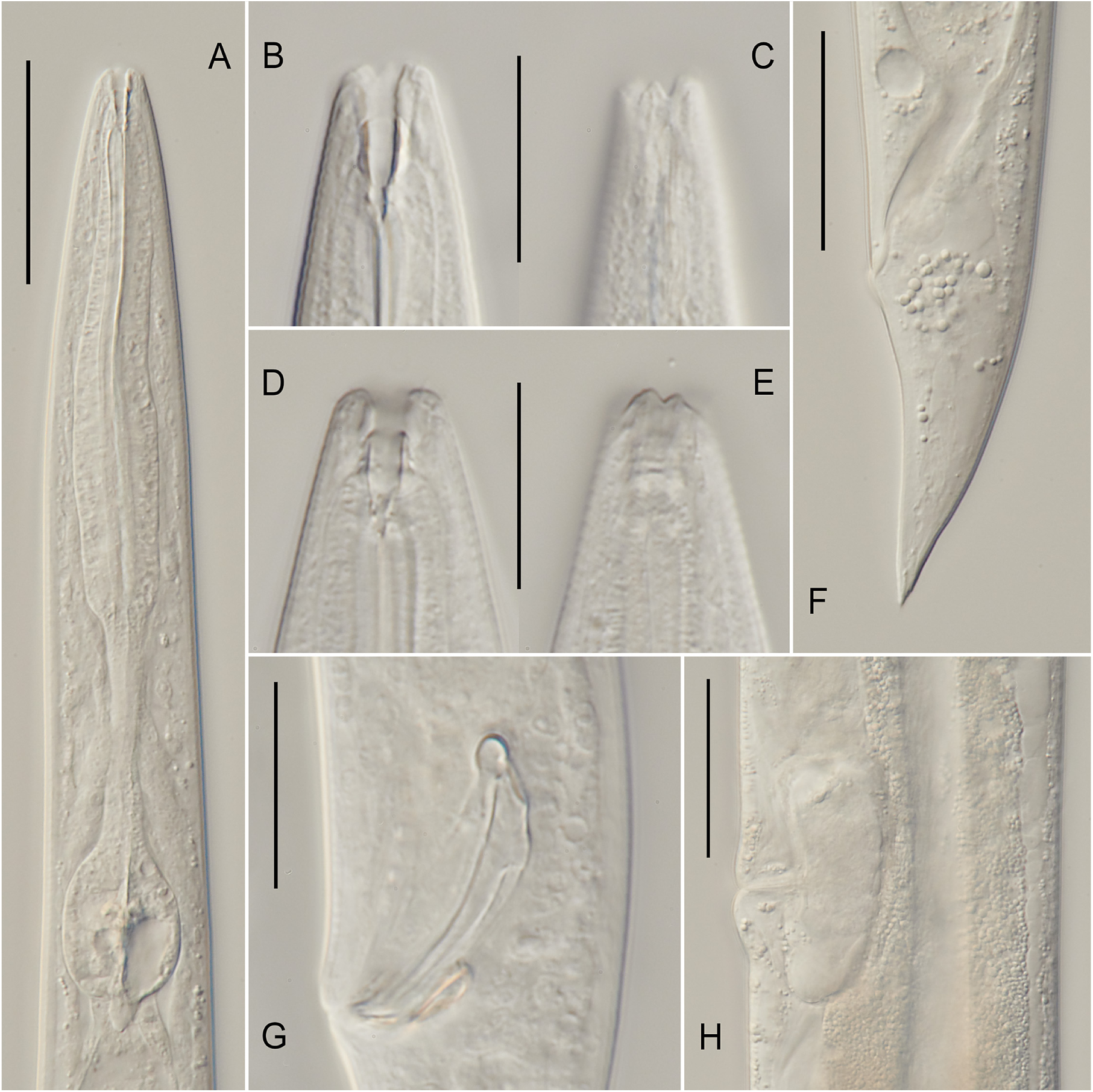
Light microscopy of ***Panagrolaimus einhardi*** sp. nov. A, pharyngeal region (ventral to the left). B, male stoma: (median section, ventral to the left). C, male labial region (surface view, ventral to the left). D, female stoma (median section, subventral). E, female labial region (surface view, subventral). F, female tail. G, male cloacal region showing spicules and gubernaculum. H, female vulval region showing vagina and post-vulval uterine sac. Scale bars: 50 µm in A, C, H; 20 µm in B-G.

**Figure 3.**
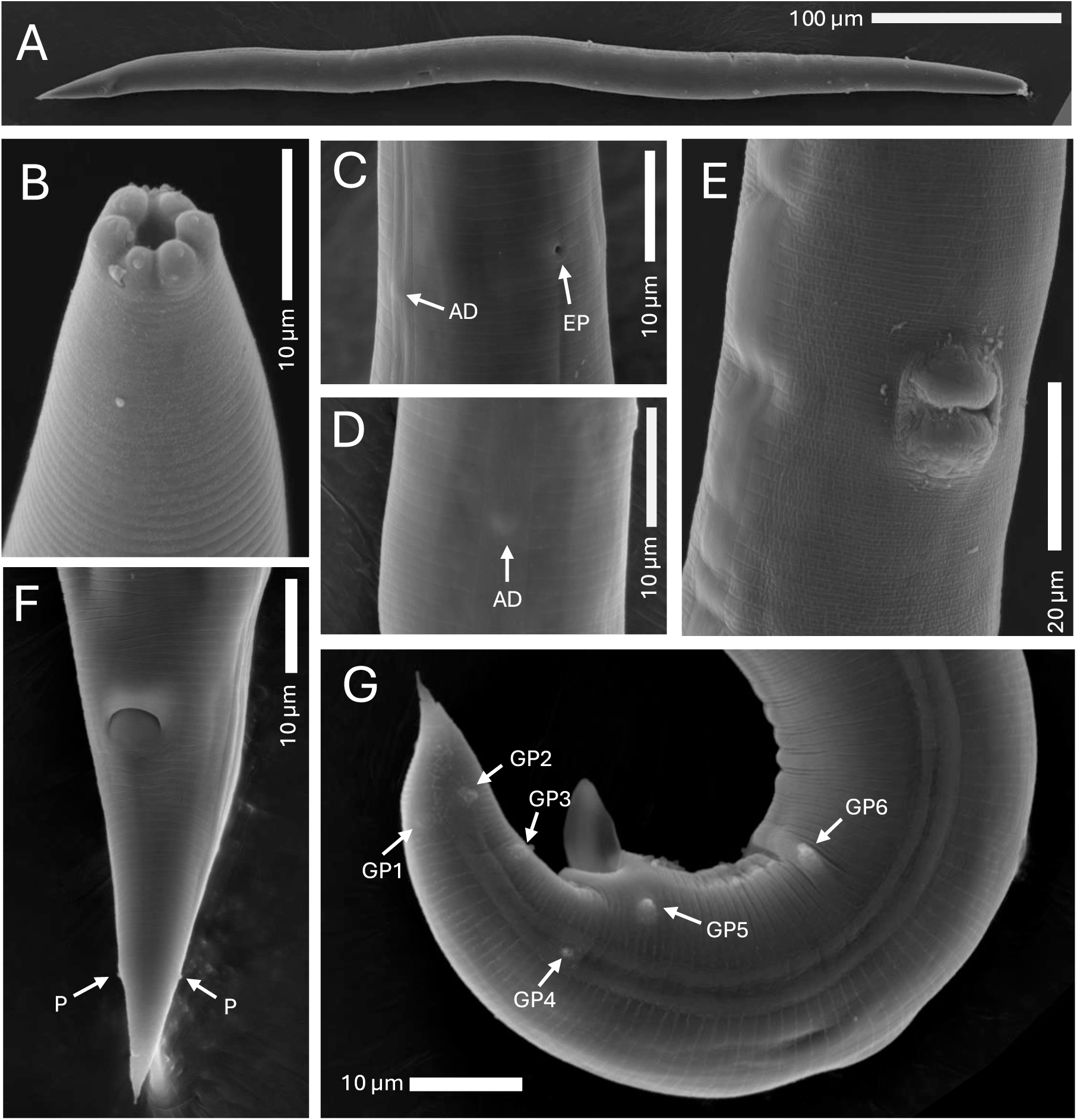
Scanning electron microscopy of ***Panagrolaimus einhardi*** sp. nov. A, full body, female. B, labial region in frontal view. C, excretory pore (EP) and anterior deirid (AD). D, lateral field, anterior deirid (AD). E, vulva. F, female tail showing anus and phasmid (P). G, male posterior end showing protruding spicules and pre-and post-cloacal genital papillae (GP).

**Table 1.** Morphometrics of *Panagrolaimus einhardi* sp. nov., *P. shuimeiren* sp. nov. and *P. nebliphilus* sp. nov. All measurements in µm and the form: mean *±* SD (range). Number of measured females and males is specified. Holotypes are separate measurements of one female each. a: body length/greatest body diameter; b: body length/neck length; c: body length/tail length, c’: tail length/tail diameter at anus or cloaca; V: % distance of vulva from anterior; Neck length: pharynx + stoma. (Comparisons to other species in Supplementary Material Figure S1-S20)

| Characters | *P. einhardi* sp. nov. |  |  | *P. shuimeiren* sp. nov. |  |  | *P. nebliphilus* sp. nov. |  |  |
| --- | --- | --- | --- | --- | --- | --- | --- | --- | --- |
|  | Holotype | 10 ♀ | 10 ♂ | Holotype | 6 ♀ | 3 ♂ | Holotype | 11 ♀ | 11 ♂ |
| Body length | 897.9 | 771.7±113.0<br>(635.4-924.2) | 737.3±97.9<br>(620.9-908.6) | 893.5 | 589.9±52.5<br>(504.1-639.3) | 609.8±11.2<br>(598.8-621.2) | 792.1 | 857.3±80.2<br>(716.1-987.2) | 752.2±62.4<br>(673.9-846.6) |
| a | 24.4 | 23.6±3.0<br>(18.6-29.8) | 25.4±3.0<br>(19.6-30.5) | 24.3 | 22.7±2.2<br>(19.2-25.6) | 23.5±1.8<br>(21.7-25.2) | 22.6 | 18.9±1.5<br>(16.2-20.5) | 25.6±1.2<br>(23.3-27.4) |
| b | 5.9 | 4.7±0.4<br>(4.2-5.4) | 4.5±0.6<br>(3.9-5.9) | 6.7 | 4.5±0.5<br>(3.6-5.0) | 4.8±0.5<br>(4.2-5.3) | 5.8 | 5.3±0.4<br>(4.7-6.0) | 5.2±0.3<br>(4.8-5.6) |
| c | 23.2 | 16.2±1.6<br>(13.9-18.7) | 18.1±1.7<br>(16.3-21.8) | 16.2 | 15.8±2.1<br>(13.0-17.9) | 15.2±1.3<br>(13.7-16.2) | 16.4 | 14.3±0.7<br>(13.5-15.6) | 14.3±1.0<br>(13.2-15.8) |
| c' | 2.2 | 2.4±0.3<br>(1.7-2.8) | 1.9±0.2<br>(1.6-2.2) | 2.2 | 2.5±0.5<br>(2.1-3.1) | 2.7±1.1<br>(2.0-3.9) | 2.3 | 2.1±0.1<br>(1.9-2.3) | 1.9±0.1<br>(1.8-2.1) |
| V (%) | 59.8 | 62.0±2.3<br>(58.3-64.9) | - | 62.5 | 59.6±3.9<br>(52.4-63.7) | - | 59.0 | 59.9±3.4<br>(51.6-63.2) | - |
| Lip region width | 7.5 | 3.5±0.4<br>(2.9-4.0) | 3.5±0.4<br>(3.0-4.0) | 9.5 | 2.9±0.2<br>(2.5-3.1) | 3.0±0.2<br>(2.8-3.1) | 9.0 | 3.6±0.3<br>(3.1-3.9) | 3.4±0.2<br>(3.1-3.8) |
| Stoma length | 9.5 | 10.5±0.4<br>(10.0-11.2) | 9.9±1.3<br>(7.6-11.5) | 10.5 | 8.9±0.7<br>(7.9-9.9) | 8.3±1.0<br>(7.5-9.4) | 9.5 | 8.7±0.2<br>(8.4-9.0) | 8.0±0.3<br>(7.6-8.4) |
| Corpus length | 87.5 | 97.6±13.4<br>(78.8-118.2) | 89.6±9.6<br>(68.6-102.9) | 77.0 | 71.0±6.9<br>(61.8-79.6) | 68.8±14.5<br>(54.7-83.6) | 80.5 | 92.8±6.2<br>(85.2-105.5) | 85.7±3.7<br>(81.7-93.8) |
| Isthmus length | 38.5 | 34.3±8.8<br>(21.6-54.6) | 38.2±5.2<br>(31.4-44.6) | 28.0 | 26.5±3.1<br>(23.5-31.3) | 27.3±4.2<br>(23.1-31.5) | 30.0 | 37.7±3.6<br>(32.8-45.4) | 34.9±2.1<br>(32.1-38.6) |
| Basal bulb | 26.5 | 23.6±1.3<br>(22.1-25.9) | 23.4±1.0<br>(22.2-25.3) | 28.0 | 24.1±0.9<br>(22.8-25.3) | 24.4±0.5<br>(23.9-24.8) | 24.5 | 24.2±2.6<br>(20.2-28.0) | 23.4±1.3<br>(20.9-25.3) |
| Pharynx | 152.5 | 155.5±14.3<br>(136.3-180.1) | 151.2±8.2<br>(133.6-162.0) | 133 | 121.6±7.2<br>(113.5-130.7) | 120.5±16.7<br>(106.5-139.0) | 136.5 | 154.7±10.5<br>(145.5-178.8) | 144.0±6.3<br>(136.9-156.0) |
| Neck length | 161.0 | 166.0±14.6<br>(146.3-191.3) | 161.0±8.3<br>(143.0-172.0) | 143.5 | 130.5±7.2<br>(122.3-139.1) | 128.8±16.2<br>(114.6-146.4) | 145.5 | 163.4±10.5<br>(154.0-187.3) | 152.0±6.5<br>(144.9-164.4) |
| Excretory pore-anterior end | 122.5 | 113.1±15.4<br>(92.3-135.0) | 115.5±11.6<br>(91.2-133.9) | - | 81.2±6.5<br>(71.9-87.4) | 80.2±11.2<br>(71.5-92.8) | 129.5 | 126.4±7.3<br>(117.8-145.1) | 119.3±5.0<br>(113.1-128.1) |
| Body width: mid-body | 37.0 | 33.1±6.2<br>(24.7-43.9) | 29.1±3.3<br>(23.4-32.7) | 37.0 | 26.3±4.3<br>(21.6-31.8) | 26.1±1.8<br>(24.1-27.6) | 35.0 | 45.5±2.4<br>(42.9-50.6) | 29.4±2.0<br>(25.8-33.0) |
| Body width: anus | 17.5 | 20.3±2.9<br>(16.7-25.4) | 21.8±1.6<br>(20.0-24.7) | 25.0 | 15.2±2.2<br>(11.6-18.2) | 16.2±4.1<br>(11.5-18.7) | 20.5 | 28.6±2.4<br>(25.7-33.9) | 28.1±1.8<br>(24.1-29.7) |
| Tail length | 38.5 | 47.8±6.3<br>(37.4-58.3) | 40.8±5.1<br>(35.2-51.2) | 55.0 | 37.5±2.6<br>(33.1-40.7) | 40.3±3.8<br>(37.0-27.6) | 48.5 | 60.1±6.6<br>(50.1-69.8) | 52.7±1.3<br>(49.9-54.1) |
| Vulva-anus | 322.0 | 281.3±64.1<br>(201.0-406.1) | - | 280.0 | 196.6±19.0<br>(174.9-217.9) | - | 276.5 | 280.9±30.8<br>(243.3-333.2) | - |
| Vulva-anterior end | - | 479.1±77.9<br>(372.5-598.8) | - | - | 352.6±47.2<br>(263.9-389.8) | - | - | 513.2±49.3<br>(418.1-590.0) | - |
| Spicule length | - | - | 27.9±7.0<br>(21.4-42.6) | - | - | 22.6±2.6<br>(19.9-25.1) | - | - | 28.3±1.8<br>(25.2-30.2) |
| Gubernaculum length | - | - | 10.7±1.8<br>(8.5-13.2) | - | - | 10.3±4.1<br>(11.5-18.7) | - | - | 11.9±0.9<br>(10.6-13.5) |

*Panagrolaimus* sp. ES5 in Schiffer et al. (2019), WormBase, ENA, NCBI GenBank, UniProt, etc

*Panagrolaimus* “bornheim” in Lewis et al. (2009)

*Panagrolaimus* sp. bornheim in Villegas et al. (2024a)

*Zoobank registration:* to be compelted upon acceptance of the manuscript

*Material examined:* Holotype female deposited in the invertebrate type collection of the Swedish Museum of Natural history, Stockholm, Sweden. Ten female and ten male paratypes on eight glass slides deposited in the Nematode Collection at the Department of Plant Pathology, Kasetsart University, Bangkok, Thailand. Additional not type specimens are deposited in the general invertebrate collection of the Swedish Museum of Natural History, Stockholm, Sweden (reg. numbers to follow). Live cultures are currently maintained in the worm∼lab at the University of Cologne and will also be deposited in the Caenorhabditis Genetics Center.

*Molecular reference data:* Nuclear genome assembly and mitochondrial genome assembly are deposited in the European Nucleotide Archive (Project number PRJEB75733).

*Type locality and habitat:* Material was collected from the soil around a dead blackberry bush (*Rubus* spp.) in Bornheim-Brenig, Germany by Prof. Einhard Schierenberg in 2006.

*Description. Female:* Body 0.63-0.92 mm long, cylindrical, ventrally curved after fixation. Both ends tapering, tail conical-elongated. Cuticle finely annulated. The lip region is rounded and continuous with the body contour. Lips high, well defined, rounded, ventrosublateral clefts are deeper than the rest; lateral lips slightly smaller than sub-dorsal and subventral ones. Stoma panagrolaimoid, with well-defined cheilostom, gymnostom, and stegostom. Cheilostom short and non-sclerotized; gymnostom long and well-sclerotized; stegostom tubular with a minute dorsal tooth. Pharynx tripartite: procorpus cylindrical, gradually widening posteriorly; isthmus narrow and tubular; basal bulb rounded and containing a well-developed valvular apparatus. Secretory-excretory pore located at the level of the isthmus, at 55-76% of neck length from the anterior end. Reproductive system monodelphic-prodelphic, ovary located on the right of the intestine. Vulva located slightly posterior to the mid-body (V = 58-65%), with protruding lips; vagina directed perpendicularly to body axis, extending inward for about 1/5-1/3 of the corresponding body diameter. Post-vulval uterine sac well developed but short, less than 12 of corresponding body diameter in length. Tail conoid, tapering to a sharp tip, often with marked narrowing in the distal half, dorsal contour convex, while ventral contour slightly concave. Phasmids are located at posterior 1/5 of the tail length.

*Male:* Similar to females in general morphology except for sexual characters, but the body is more curved ventrally, forming a J-shape upon fixation. Reproductive system monorchic, testis reflexed ventrally. Tail conical, distally curved ventrally. Spicules ventrally curved, 21-42 µm in length. Gubernaculum deltoid, 8-13 µm long. Six pre-and post-cloacal genital papillae (GP) arranged as depicted on Fig. 3G.

*Etymology: Panagrolaimus einhardi* sp. nov. is named in honour of eminent nematologist and developmental biologist Prof. Dr. Einhard Schierenberg. Prof. Schierenberg dedicated his work-life to the study of roundworms and was one of the first, who extended his scope of research away from the model system *C. elegans* to evolutionary distant species across the phylum Nematoda. This led him and his team to important discoveries on the variation of early development in nematodes. Einhard was a very respected, cherished, and universally liked member of the worm community, who is duly missed. Naming this species, which he himself sampled near Cologne in Germany, after him, we hope to uphold the memory of an incredibly curious scientist and a great friend and supervisor.

*Diagnosis: Panagrolaimus einhardi* sp. nov. is characterized by a moderately sized body, measuring 0.63-0.92 mm in females and 0.62-0.90 mm in males. Labial region with well developed separate lips, of which lateral ones are smaller. Stoma panagrolaimoid, consisting of a short cheilostom, a well-developed and cuticularized gymnostom, and a narrow stegostom with minute internal structures. Pharynx with a cylindrical procorpus and a weakly swollen metacorpus; nerve ring and excretory pore located at the level of the isthmus. Female reproductive system monodelphic-prodelphic, with the vulva situated slightly posterior to mid-body (V = 58-64 %) and bearing prominently protruding lips. Vagina transverse and relatively short. Female tail conoid, tapering to an acute terminus. Male tail conoid, curved ventrally, with spicules moderately curved ventrad (21-42 µm long) and a deltoid gubernaculum (8-13 µm), typically less than half the spicule length.

*Relationships:* The new species can be distinguished from related, similar species by the following features:From *P. shuimeiren* sp. nov. by a differently shaped lip region with lateral lips smaller then the rest (*vs* lips equal in size and partially amalgamated in pairs), generally longer body length (635.4-924.2 µm vs. 504.1-639.3 µm), also seen in the distance of excretory pore to anterior end (92.3-135.0 µm vs. 71.9-87.4 µm) together with a wider body at anus level (16.7-25.4 µm vs. 11.6-18.2 µm) and the absence of a midventral papillae (MP).

From *P. nebliphilus* sp. nov. by a differently shaped lip region with lateral lips smaller then the rest (*vs* lips equal in size and amalgamated in pairs), stoma without distinct dorsal tooth (*vs* with distinct dorsal tooth), more narrow body width (mid: 24.7-43.9 µm; anus: 16.7-25.4 µm vs. mid: 42.9-50.6 µm; anus: 25.7-33.9 µm) and the number of genital papillae in males (GP; six vs. eight pairs).

From *P. superbus* (after Mehdizadeh et al. (2013)) by a differently shaped lip region with lateral lips smaller then the rest (*vs* lips equal in size and amalgamated in pairs), phasmid located at posterior 1/5th of tail length in females (*vs* at middle of tail), shorter spicule (20.9-35.0 µm vs. 35-37 µm) and gubernaculum (8.9-12.4 µm vs. 13-15 µm) in males and a generally narrower body width, as well as different basal bulb sizes (22.3-25.0 µm vs. 25.1-33.3 µm), stoma length (10.1-10.9 µm vs. 25.1-33.3 µm), and the ratio of body length to tail length (c) (14.6-17.8 vs. 19.0-21.8) and the ratio of tail length to tail diameter (c’) (2.1-2.7 vs. 1.7-1.9).

From *P. davidi* (after Timm (1971)) by a stoma without distinct dorsal tooth (*vs* with distinct dorsal tooth), smaller basal bulb (22.3-25.0 µm vs. 26-29 µm), as well as a different number of genital papillae (GP; six vs. eight pairs). *P. davidi* is an endemic species to continental Antarctica and has not been recorded elsewhere (except as a laboratory strain).

From *P. namibiensis* (after Rawson et al. (2024)) by a differently shaped lip region with lateral lips smaller then the rest (*vs* lips equal in size and amalgamated in pairs), stoma without distinct dorsal tooth (*vs* with small dorsal tooth),smaller ratio of body length to tail length (c) (14.6-17.8 vs. 20.2-22.8), a larger ratio of tail length to tail diameter (c’) (2.1-2.7 vs. 1.6-2), and a smaller ratio of body length to neck length (b) (4.3-5.1 vs. 5.8-6.6), as well as a different number of genital papillae (GP; six vs. seven pairs).

From *P. kolymaensis* (after Shatilovich et al. (2023)) specifically by gonochoristic reproduction mode indicated by the abundance of males.

From *P. subelongatus* (after Mehdizadeh et al. (2013)) by a differently shaped lip region with lateral lips smaller then the rest (*vs* lips fully amalgamated in pairs), smaller ratio of body length to tail length (c) (14.6-17.8 vs. 19.4-21.6), ratio of tail length to tail diameter (c’) (2.1-2.7 vs 1.6-2.0), and the distance of excretory pore to anterior end (97.7-128.6 µm vs. 137.7-144.5 µm).

From *P. rigidus* (after Seddiqi et al. (2016)) by a differently shaped lip region with lateral lips smaller then the rest (*vs* lips amalgamated in pairs), conoid tail (*vs* with lanceolate tip), smaller ratio of body length to neck length (b) (4.3-5.1 vs. 6.0-6.4), a larger ratio of body length to tail length (c) (14.6-17.8 vs. 13.6-14.0), a smaller ratio of tail length to tail diameter (c’) (2.1-2.7 vs. 3.1-3.7) and a different corpus length (84.2-111.0 µm vs. 73.5-78.3 µm), as well as the proportional distance from vulva to anterior end (V) (59.7-64.2 % vs. 56.7-59.7 %).

From *P. orientalis* (after Seddiqi et al. (2016)) by a differently shaped lip region with lateral lips smaller then the rest (*vs* lips equal in size and amalgamated in pairs), stoma without distinct dorsal tooth (*vs* with small dorsal tooth), phasmid located at posterior 1/5th of tail length in females (*vs* at middle of tail), larger ratio of body length to body diameter (a) (20.6-26.6 vs. 16.2-19.6), larger ratio of body length to tail length (c) (14.6-17.8 vs. 10.7-12.3), a larger ratio of tail length to tail diameter (c’) (2.1-2.7 vs. 2.9-3.5), and larger corpus length (84.2-111.0 µm vs. 71.9-78.3 µm), as well as proportional distance from vulva to anterior end (V) (59.7-64.2 % vs. 55.9-59.5 %). Original description of *Panagrolaimus orientalis* (Korentchenko, 1986) specifies an anisomorphic stoma with two subdorsal teeth – condition that is different from both *P. einhardi* sp. nov. and Seddigi’s redescription.

From *P. facetus* (after Mehdizadeh et al. (2013)) by a differently shaped lip region with lateral lips smaller then the rest (*vs* lips equal in size and amalgamated in pairs), phasmid located at posterior 1/5th of tail length in females (*vs* at anterior 1/3rd of tail), larger ratio of body length to tail length (c) (14.6-17.8 vs. 12.5-13.5), with a different stoma length (10.1-10.9 vs. 12.1-13.1), as well as a proportionally larger distance from vulva to anterior end (V) (59.7-64.2 % vs. 56.9-59.3 %).

From *P. labiatus* (after Yadav et al. (2022)) by a differently shaped lip region with lateral lips smaller then the rest (*vs* lips equal in size and amalgamated in pairs), phasmid located at posterior 1/5th of tail length in females (*vs* at anterior 1/3rd of tail), larger ratio of body length to tail length (c) (14.6-17.8 vs. 11.2-13.4), smaller ratio of tail length to tail diameter (c’) (2.1-2.7 vs. 2.9-3.7), as well as larger proportional distance from vulva to anterior end (V) (59.7-64.2 % vs. 55.6-58.4 %).

From *P. concolor* (after Mehdizadeh et al. (2013)) by a differently shaped lip region with lateral lips smaller then the rest (*vs* lips equal in size and amalgamated in pairs), stoma without dorsal tooth (*vs* with small dorsal tooth), phasmid located at posterior 1/5th of tail length in females (*vs* at middle of tail), smaller ratio of body length to neck length (b) (4.3-5.1 vs. 5.1-7.3), shorter distance of excretory pore to anterior end (97.7-128.6 µm vs. 149.2-161.8 µm), slimmer body width (mid-body: 26.9-39.4 µm vs. 49.0-52.6 µm; anus: 17.5-23.2 µm vs. 25.7-29.1 µm), and smaller basal bulb size (22.3-25.0 µm vs. 26.5-26.5 µm), as well as shorter tail length (41.5-54.1 µm vs. 57.9-75.1 µm).

From *P. papillosus* (after Mehdizadeh et al. (2013)) by a smaller ratio of body length to tail length (c) (14.6-17.8 vs. 19.2-20.4), larger ratio of tail length to tail diameter (c’) (2.1-2.7 vs. 1.7-1.9), with a different stoma length (10.1-10.9 µm vs. 11.4-13.2 µm), basal bulb size (22.3-25.0 µm vs. 28.3-30.5 µm) and tail length (41.5-54.1 µm vs. 37.9-40.1 µm). Type population of *P. papillosus* described by Loof is most likely asexual (Loof, 1971) while both *P. einhardi* and Mehdizadeh’s population are gonochoristic.

From *P. leperisini* (after Massey (1974); Girgan et al. (2018)) by a papilliform anterior sensilla (*vs* setiform), stoma without distinct dorsal tooth (*vs* with distinct dorsal tooth), smaller ratio of body length to body diameter (a) (20.6-26.6 vs. 29.5-34.9), by a smaller ratio of body length to tail length (c) (14.6-17.8 vs. 18.9-19.2), as well as larger proportional distance from vulva to anterior end (V) (59.7-64.2 % vs. 55.9-56.8 %) and the number of genital papillae in males (six vs. five pair)

> Phylum Nematoda Potts, 1932
>
> Class Chromadorea Inglis, 1983
>
> Suborder Tylenchina Thorne, 1949
>
> Family Panagrolaimidae Thorne, 1937
>
> **Genus *Panagrolamus*** Fuchs, 1930
>
> ***Panagrolaimus shuimeiren* sp. nov.** (Figures 1C-D, 4-5; Table 1)

**Figure 4.**
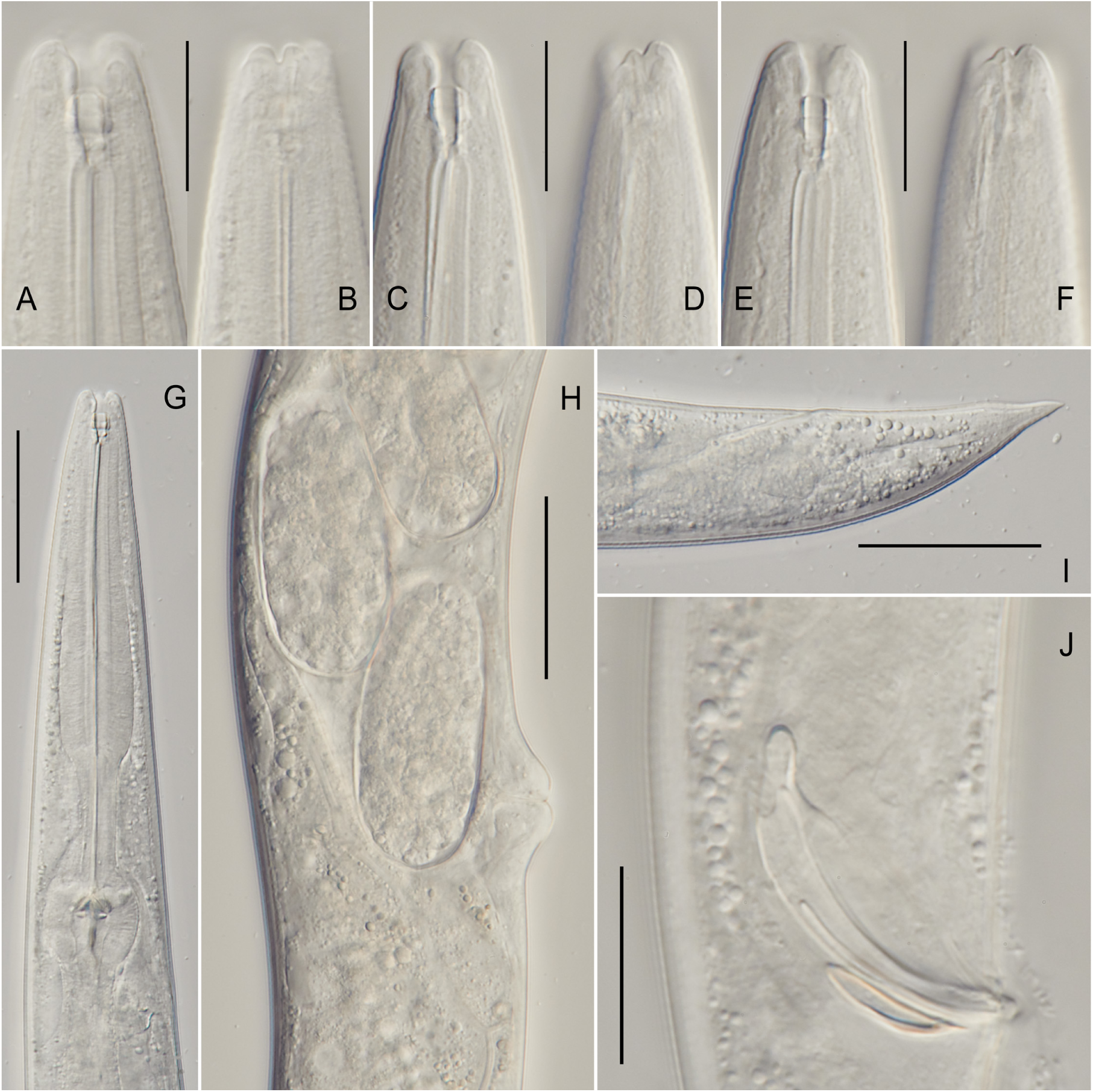
Light microscopy of ***Panagrolaimus shuimeiren*** sp. nov. A, female stoma (median section, ventral to the right). B: female labial region (surface view, ventral to the right). C, E, male stoma (median section, ventral to the right). D, F, male labial region (surface view, ventral to the right). G, pharyngeal region (ventral to the right). H, female vulval region, showing intruterine eggs, vagina and post-vulval uterine sac. I, female tail. J, male cloacal region showing spicules and gubernaculum. Scale bars: 20 µm in A-F and J; 50 µm in G-I.

**Figure 5.**
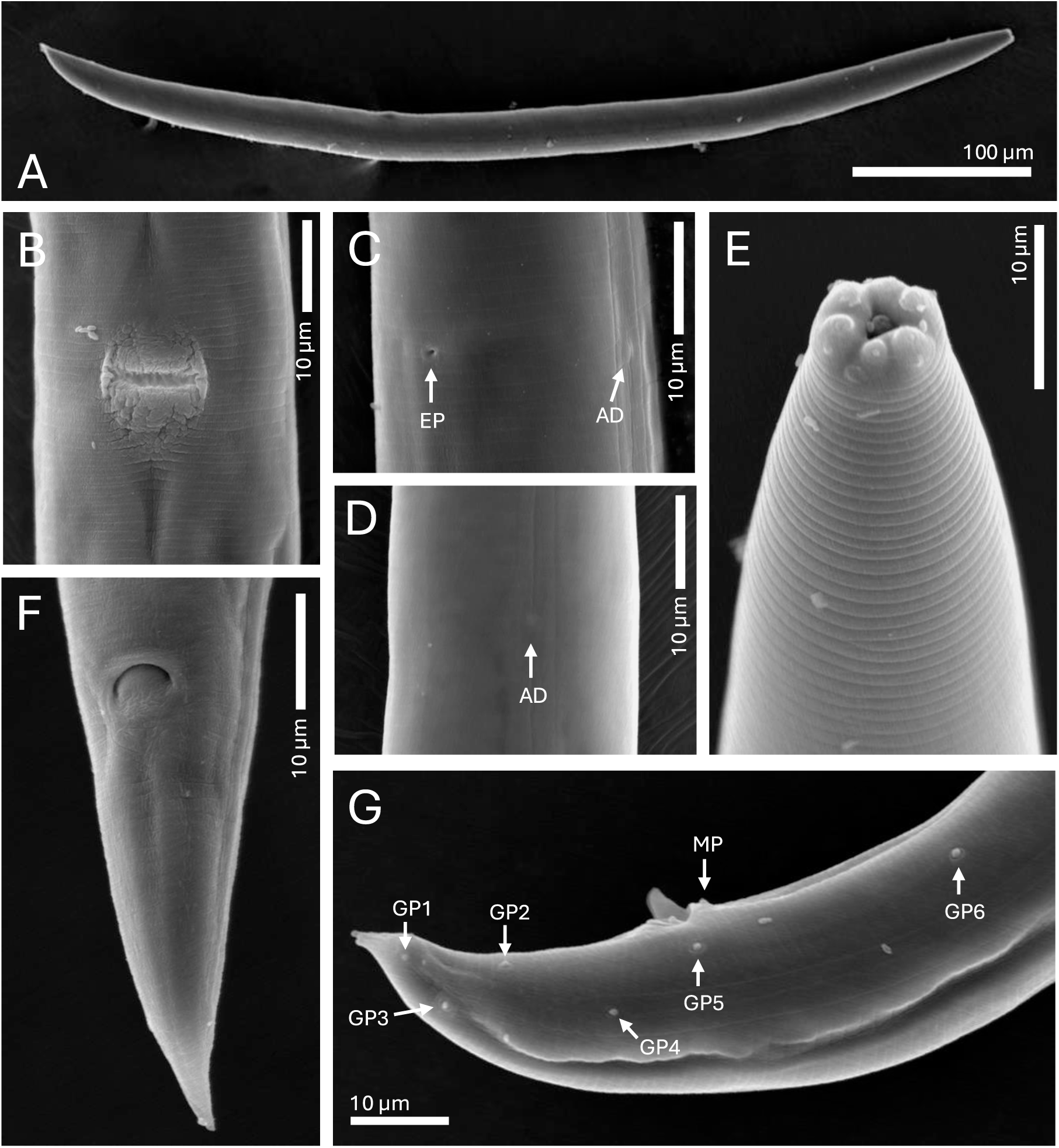
Scanning electron microscopy of ***Panagrolaimus shuimeiren*** sp. nov. A, full body, female. B, vulva. C, lateral field, excretory pore (EP) and anterior deirid (AD). D, anterior deirid (AD). E, labial region in frontal view. F, female tail showing anus. G, male posterior end, showing protruding spicules, pre- and post-cloacal genital papillae (GP), midventral papilla (MP).

#### *Zoobank registration:* to be completed upon acceptance of the manuscript

*Type material:* Holotype female deposited in the invertebrate type collection of the Swedish Museum of Natural history, Stockholm, Sweden. Six female and three male paratypes on four glass slides deposited at the Institute for Zoology, University of Cologne, Germany. Additional not type specimens are deposited in the general invertebrate collection of the Swedish Museum of Natural History, Stockholm, Sweden (reg. numbers to follow). Live cultures are currently maintained in the worm∼lab at the University of Cologne and will also be deposited in the Caenorhabditis Genetics Center.

*Molecular reference data:* Nuclear genome assembly will be deposited in the European Nucleotide Archive (Project number PRJEB75733).

*Type locality and habitat:* Specimens were recovered from arid soil in the Namib Desert, Skeleton Coast area, Namibia (20°16’57.9”S, 13°40’58.2”E), in a depth of 5-10cm close to *Kissenia* spp. at 520 m.a.s.l.

*Description. Female:* Body length 0.5-0.63 mm, spindle-shaped, nearly straight after fixation. Cuticle finely annulated. The lip region is continuous with the body contour. Lips high, well defined, prounded, partially amalgamated at their bases into three pairs. Stoma panagrolaimoid, its total length ≤ 10 µm, maximum stoma width ≤ 3 µm, composed of a short cheilostom, a wide and relatively long gymnostom with strongly sclerotized rhabdia, and a gradually tapering stegostom without distinct dorsal denticle. Pharynx tripartite: procorpus cylindrical, gradually widening posteriorly; isthmus narrow and tubular; basal bulb rounded and containing a well-developed valvular apparatus. Secretory-excretory pore located at the level of the isthmus. Reproductive system monodelphic-prodelphic, situated dorsally and to the right of the intestine. Vulva positioned slightly posterior to the mid-body (V= 52-63 %; mean 59 %), with distinctly protruding lips; vagina transverse, directed perpendicularly to body axis, extending inward less than 1/4 of the corresponding body width. Tail elongate-conoid to conoid, tapering to a pointed terminus, with distinct terminal section which is often bent dorsally. Phasmids are located at posterior 1/3 of the tail length.

*Male:* Body length 0.59-0.62 mm; similar to females in general morphology except for sexual characters, but with the posterior body region more strongly curved ventrally (J-shaped after fixation). Reproductive system monorchic; testis reflexed ventrally. Tail conical and curved ventrad distally, with distinct digitate terminal part about 1/5 of total tail length. Spicules ventrally curved, measuring 19-25 µm in length. Gubernaculum well developed, measuring 11-18 µm, typically less than half the spicule length. Six pre-and post-cloacal genital papillae (GP) and one midventral papilla (MP), arranged as depicted on Fig. 5G.

*Etymology: P. shuimeiren* sp. nov. is named for its ability to reduce metabolic activity and enter a cryptobiotic state. Its epithet “shuimeiren,” from the Mandarin “shùı m_̌_ei ŕen,” means “Sleeping Beauty,” reflecting its prolonged dormant state akin to the fairytale character’s deep sleep.

*Diagnosis: Panagrolaimus shuimeiren* sp. nov is characterized by a moderately small, spindle-shaped body (females: 0.50-0.63 mm; males: 0.59-0.62 mm). Labial region with well developed lips, partially amalgamated at their bases into three pairs. Stoma panagrolaimoid, with a heavily sclerotized gymnostom, narrow stegostom without defined dorsal denticle. Pharynx with cylindrical procorpus, narrow isthmus, and a strongly muscular basal bulb. Excretory pore is located at the isthmus level. Female reproductive system monodelphic-prodelphic, with a transverse vagina and vulva positioned slightly posterior to mid-body (V = 52-63%), showing prominent, protruding lips. Female tail elongate-conoid, tapering smoothly to a fine, pointed terminus, often dorsally curved. Males are distinguished by their more ventrally curved posterior body, conical tail, moderately long spicules (19-25 µm), and a well-developed gubernaculum measuring 11-18 µm, typically less than half the spicule length.

*Relationships:* The new species can be distinguished from related, similar species by the following features: From *P. nebliphilus* by a differently shaped lip region with lips only partially amalgamated into pairs (*vs* lips nearly completely amalgamated in pairs), stoma without distinct dorsal tooth (*vs* with distinct dorsal tooth), narrower body width (mid: 22.0-30.5 µm vs. 43.1-47.9 µm; anus: 13.0-17.4 µm vs. 26.2-31.0 µm), different corpus length (64.2-77.9 µm vs. 86.6-99.0 µm), a larger ratio of body length to the greatest body diameter (a) (20.6-24.9 µm vs. 17.4-20.3 µm), and the distance of excretory pore to anterior end (74.7-87.6 µm vs. 119.1-133.8 µm), as well as different spicule lengths in males (20.0-25.2 µm vs. 26.5-30.1 µm).

From *P. superbus* (after Mehdizadeh et al. (2013)) by a differently shaped lip region with lips only partially amalgamated into pairs (*vs* lips nearly completely amalgamated in pairs), phasmid located at posterior 1/3rd of tail length in females (*vs* at middle of tail), smaller ratio of body length to tail length (c) (13.8-17.9 vs. 19.0-21.8), larger ratio of tail length to tail diameter (c’) (2.1-3.0 vs. 1.7-1.9), with a shorter corpus length (64.2-77.9 µm vs. 95.7-100.5 µm), and the distance of excretory pore to anterior end (74.7-87.6 µm vs. 148.3-156.1 µ), as well as different spicule lengths in males (20.0-25.2 µm vs. 35.0-37 µm).

From *P. davidi* (after Timm (1971)) by a differently shaped lip region with all lips equal in size (vs lateral lips, called “submedian lobe” in the original description, being smaller), stoma without distinct dorsal tooth (*vs* with distinct dorsal tooth), shorter body length (537.3-642.4 µm vs. 850.0-970.0 µm), shorter basal bulb size (23.2-24.9 µm vs. 26.0-29.0 µm), as well as a different number of genital papillae. *P. davidi* is an endemic species to continental Antarctica and has not been recorded elsewhere (except as a laboratory strain).

The new species is very similar to *P. namibiensis*, which was recently described from Namib desert. However, it differs from *P. namibiensis* (after Rawson et al. (2024)) by a smaller ratio of body length to neck length (b) (4.0-5.0 vs. 5.8-6.6), ratio of body length to tail length (c) (13.8-17.9 vs. 20.2-22.8), ratio of tail length to tail diameter (c’) (2.1-3.0 vs. 1.6-2.0), with a different corpus length (64.2-77.9 µm vs.84.9-91.7 µm), as well as different spicule length in males (20.0-25.2 µm vs. 27.3-31.3 µm). 18S rDNA sequences of *P. namibiensis* were not included in the current phylogeny due to them being less than 600 bases long, but in the molecular phylogeny from the original description this species clusters with *P. kolymaensis*, and thus is also genetically close to *P. shuimeiren* sp. nov., nonetheless the overlapping sections of 18S rDNA from both species (558 bp) differ in 28 positions.

From *P. kolymaensis* (after Shatilovich et al. (2023)) specifically by gonochoristic reproduction mode indicated by the abundance of males.

From *P. subelongatus* (after Mehdizadeh et al. (2013)) by a differently shaped lip region with all lips equal in size (*vs* lips completely amalgamated in pairs), smaller ratio of body length to tail length (c) (13.8-17.9 vs. 19.4-21.6), larger ratio of tail length to tail diameter (c’) (2.1-3.0 vs. 1.6-2.0), with a shorter corpus length and shorter distance excretory pore to anterior end (74.7-87.6 µm vs. 137.7-144.5 µm), as well as different spicule length in males (20.0-25.2 µm vs. 29.0-30.0 µm).

From *P. rigidus* (after Seddiqi et al. (2016)) by a conoid tail (*vs* with lanceolate tip), ratio of body length to neck length (b) (4.0-5.0 vs. 6.0-6.4), a smaller ratio of tail length to tail diameter (c’) (2.1-3.0 vs. 3.1-3.7), with a different distance from excretory pore to anterior end (74.7-87.6 µm vs. 126.7-131.3 µm), as well as different spicule length in males (74.7-87.6 µm vs. 126.7-131.3 µm).

From *P. orientalis* (after Seddiqi et al. (2016)) by a stoma without distinct dorsal tooth (*vs* with small dorsal tooth), phasmid located at posterior 1/3rd of tail length in females (*vs* at middle of tail), larger ratio of body length to greatest diameter (a) (20.6-24.7 vs. 16.2-19.6), a larger ratio of body length to tail length (c) (20.6-24.9 vs. 16.2-19.6), with a smaller distance of excretory pore to anterior end (74.7-87.6 µm vs. 103.4-127.0 µm). Original description of *Panagrolaimus orientalis* (Korentchenko, 1986) specifies an anisomorphic stoma with two subdorsal teeth – condition that is different from both *P. shuimeiren* sp. nov. and Seddigi’s redescription.

From *P. facetus* (after Mehdizadeh et al. (2013)) by a phasmid located at posterior 1/3rd of tail length in females (*vs* at anterior 1/3rd of tail), ratio of body length to tail length (c) (13.8-17.9 vs. 12.5-13.5), with a different corpus length (64.2-77.9 µm vs. 94.9-109.5 µm), as well as distance of excretory pore to anterior end (74.7-87.6 µm vs. 111.7-159.9 µm).

From *P. labiatus* (after Yadav et al. (2022)) by a phasmid located at posterior 1/3rd of tail length in females (*vs* at anterior 1/3rd of tail), larger ratio of body length to tail length (c) (13.8-17.9 vs. 11.2-13.4), and more narrow body width (mid: 22.0-30.5 µm vs. 32.0-40.0 µm; anus: 13.0-17.4 µm vs. 17.7-24.3 µm), as well distance of excretory pore to anterior end (74.7-87.6 µm vs. 107.4-124.6 µm).

From *P. concolor* (after Mehdizadeh et al. (2013)) by a stoma without dorsal tooth (*vs* with small dorsal tooth), smaller ratio of body length to neck length (b) (4.0-5.0 vs. 5.1-7.3), a more narrow body width (mid: 22.0-30.5 µm vs. 49.0-52.6 µm; anus: 13.0-17.4 µm vs. 25.7-29.1 µm), a shorter corpus length (64.2-77.9 µm vs. 78.4-84.8 µm) and shorter distance from excretory pore to anterior end (74.7-87.6 µm vs. 149.2-161.8 µm).

From *P. papillosus* (after Mehdizadeh et al. (2013)) by a smaller ratio of body length to tail length (c) (13.8-17.9 vs. 19.2-20.4), ratio of tail length to tail diameter (c’) (2.1-3.0 vs. 1.7-1.9), with a different corpus length (64.2-77.9 µm vs. 95.8-98.6 µm) and distance from excretory pore to anterior end (74.7-87.6 µm vs. 133.0-143.2 µm). Type population of *P. papillosus* described by Loof is most likely asexual (Loof, 1971) while both *P. shuimeiren* and Mehdizadeh’s population are gonochoristic

From *P. leperisini* (after Massey (1974); Girgan et al. (2018)) by a papilliform anterior sensilla (*vs* setiform), stoma without distinct dorsal tooth (*vs* with distinct dorsal tooth), smaller ratio of body length to greatest diameter (a) (20.6-24.9 vs. 29.5-34.9), a smaller ratio of body length to tail length (c) (13.8-17.9 vs. 18.9-19.2), narrower body width (mid: 22.0-30.5 µm vs. 31.0-33.0 µm; anus: 13.0-17.4 µm vs. 18.0-21.0 µm), a shorter corpus length (64.2-77.9 µm vs. 107.0-114.0 µm), and shorter spicule length (20.0-25.2 µm vs. 28.0-29.0 µm).

> Phylum Nematoda Potts, 1932
>
> Class Chromadorea Inglis, 1983
>
> Suborder Tylenchina Thorne, 1949
>
> Family Panagrolaimidae Thorne, 1937
>
> Genus *Panagrolaimus* Fuchs, 1930
>
> Panagrolaimus nebliphilus sp. nov.
>
> (Figures 1E-F, 6-7; Table 1)

**Figure 6.**
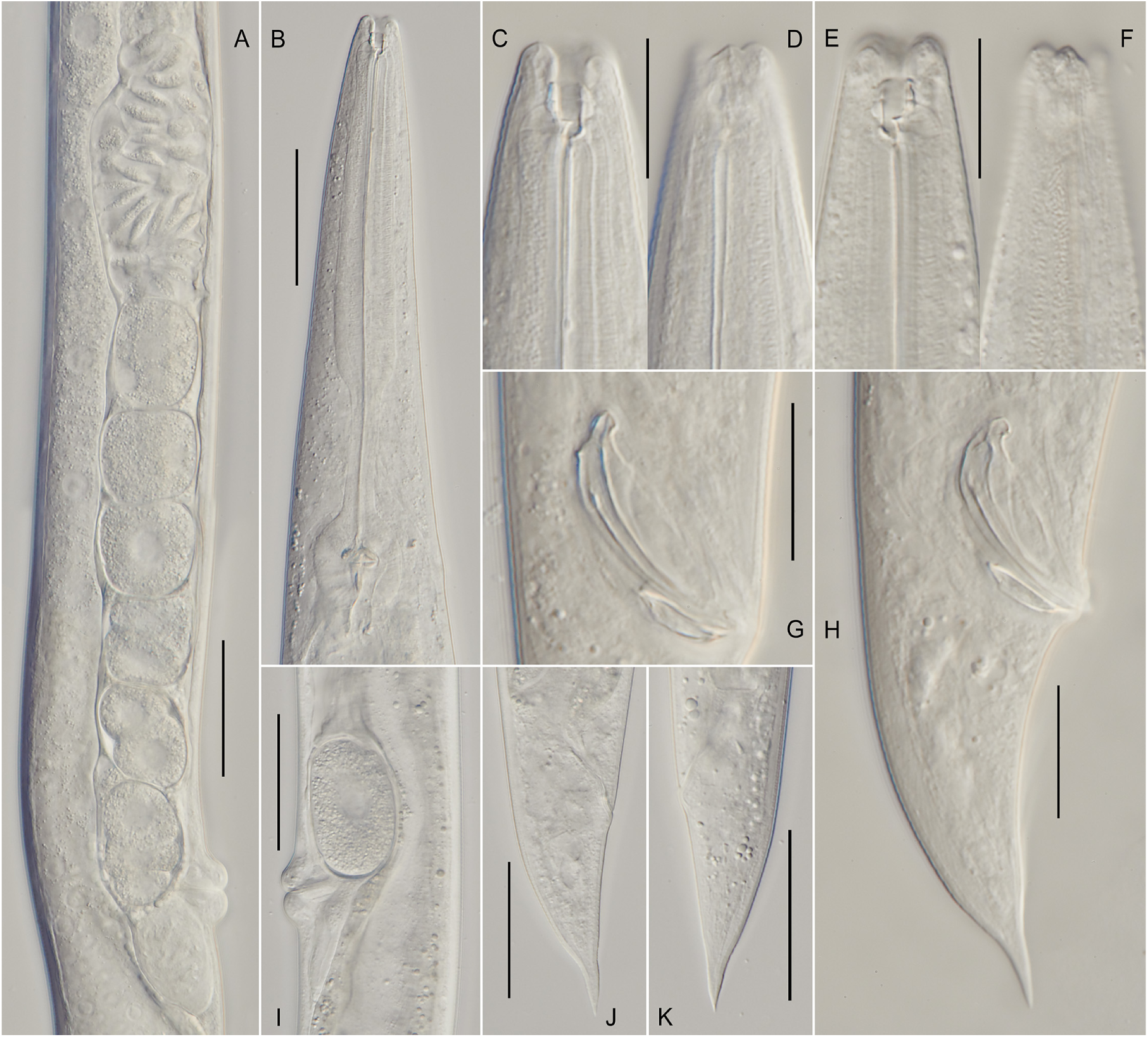
Light microscopy of ***Panagrolaimus nebliphilus*** sp. nov. A, female vulval region showing spermatheca, uterus filled with egg, vagina and post-vulval uterine sac. B, pharyngeal region, ventral to the right. C, female stoma (median section, ventral to the right). D, female labial region (surface view, ventral to the right). E, female stoma (median section, ventral to the left). F, female labial region (surface view, ventral to the left). G-H, male caudal region showing spicules and gubernaculum. I, female vulval region showing intrauterine egg, vagina and post-vulval uterine sac. J-K, female tail. Scale bars: 50 µm in A-B and I-K; 20 µm in C-H.

**Figure 7.**
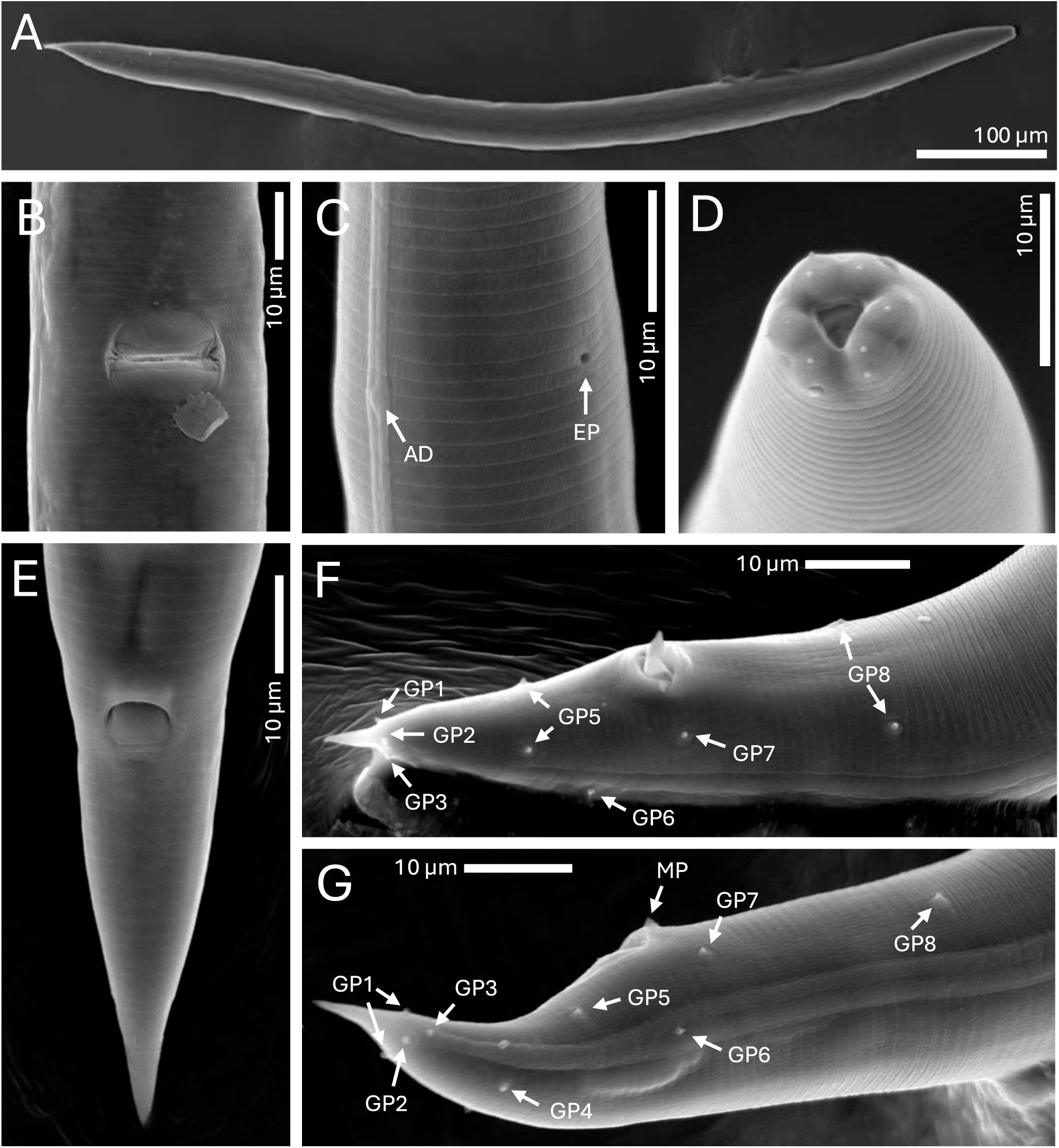
Scanning electron microscopy of ***Panagrolaimus nebliphilus*** sp. nov. A, full body, female. B, vulva. C, excretory pore (EP) and anterior deirid (AD). D, labial region in frontal view. E: female tail showing anus. F-G, male posterior end showing protruding spicules, pre- and post-cloacal genital papillae (GP), midventral papilla (MP).

Syn.: *Panagrolaimus* sp. PAP.22.29 in Villegas et al. (2025)

*Zoobank registration:* to be completed upon acceptance of the manuscript

*Type material:* Holotype female deposited in the invertebrate type collection of the Swedish Museum of Natural history, Stockholm, Sweden. Eighteen specimens of each sex, mounted on eight glass slides, are deposited in the Nematode Collection of the Department of Plant Pathology, Kasetsart University, Bangkok, Thailand. Additional not type specimens are deposited in the general invertebrate collection of the Swedish Museum of Natural History, Stockholm, Sweden (reg. numbers to follow). Live cultures are currently maintained in the worm∼lab at the University of Cologne and will also be deposited in the Caenorhabditis Genetics Center.

*Molecular reference data:* Nuclear genome assembly will be deposited in the European Nucleotide Archive (Project number PRJEB75733).

*Type locality and habitat:* Specimens were collected from saline-rich, xeric soil in the Atacama Desert, Paposo, Chile (25°14’6”S, 70°25’31”W), at a depth of 5-10 cm close to *Solanum* spp. at 220 m.a.s.l.

*Description. Female:* Body length 0.72-0.99 mm, cylindrical, slightly ventrally curved after fixation. Cuticle finely annulated. The lip region is continuous with the adjoining body contour. Lips low, rounded, nearly completely amalgamated at their bases. Stoma panagrolaimoid with distinct and separate cheilostom, gymnostom, and stegostom. Cheilostom short, with weakly refringent rhabdia; gymnostom shorter than cheilostom, with cuticularized with rhabdia, triangular in transverse section; stegostom broad, anisomorphic, with heavily cuticularised prostegorhabdia, and relatively large dorsal metastegostomal tooth. Muscular tissue is more strongly developed on the dorsal side of the stoma. Pharynx with procorpus cylindrical and longer than the metacorpus. Metacorpus slightly swollen; isthmus narrow and well-separated from the metacorpus. Basal bulb ovoid to spheroid, containing a well-developed valvular apparatus. Secretory-excretory pore located at 73-80 % of neck length, at the isthmus level. Reproductive system monodelphic-prodelphic, situated on the right of the intestine. Vulva located slightly posterior to mid-body (V = 51-63%), with distinctly protruding lips.; vagina transverse, directed perpendicularly to body axis, extending inward less than 1/4 of the corresponding body width. Tail conical-elongate, with digitate terminal part tapering to a finely pointed terminus, often slightly bent dorsad. Phasmids are located at posterior 1/3 of the tail length.

*Male:* General morphology similar to females, but with a more strongly ventrally curved posterior region (J-shaped after fixation). Reproductive system monorchic; testis reflexed dorsally in the anterior body region. Tail conical, ventrally curved at the distal end, with distinct digitate terminal part about 1/5 of total tail length. Spicules curved ventrally, slender, and pointed at the tip, measuring 25-30 µm in length. Gubernaculum well-developed, deltoid in shape, measuring up to 13 µm in length, typically less than half the spicule length. Eight pre-and post-cloacal genital papillae (GP) and one midventral papilla (MP), arranged as depicted on Fig. 7F-G.

*Etymology: P. nebliphilus* is named after the special fog event in the coastal region around Paposo, Chile (i.e. the camanchaca), where nebula = fog and -philus = loving, translating to the “fog-loving” *Panagrolaimus*.

*Diagnosis. Panagrolaimus nebliphilus* sp. nov is characterized by a medium body size (females: 0.72-0.99 mm). Labial region with six low lips amalgamater into three pairs. Stoma panagrolaimoid with cuticularized anisomorphic gymnostom and a relatively large dorsal tooth. Pharynx with slightly swollen metacorpus. Excretory pore positioned at the isthmus level. The species exhibits a vulva positioned posterior to mid-body (V = 51-63 %) with protruding lips, and a conical-elongate tail terminating, with distinct digitate terminal part. Males are distinguished by a dorsally reflexed testis, a ventrally curved conical tail, spicules 25-30 µm long, and a short, deltoid gubernaculum ≤ 13 µm.

*Relationships:* The new species can be distinguished from related, similar species by the following features:

From *P. superbus* (after Mehdizadeh et al. (2013)) by a stoma with distinct dorsal tooth (*vs* without), phasmid located at posterior 1/3rd of tail length in females (*vs* at middle of tail), smaller ratio of body length to tail length (c) (13.6-15.0 vs. 19.0-21.8), a smaller ratio of tail length to tail diameter (c’) (2.0-2.2 vs. 1.7-1.9), as well as different spicule (26.5-30.1 µm vs. 35.0-37.0 µm) and gubernaculum length in males (11.0-12.8 µm vs. 13.0-15.0 µm).

From *P. davidi* (after Timm (1971)) by a differently shaped lip region with all lips equal in size (vs lateral lips, called “submedian lobe” in the original description, being smaller), larger ratio of body length to neck length (b) (4.8-5.7 vs. 4.2-4.8) and smaller ratio of body length to tail length (c) (13.6-15.0 vs. 17.1-21.3). *P. davidi* is an endemic species to continental Antarctica and has not been recorded elsewhere (except as a laboratory strain).

From *P. namibiensis* (after Rawson et al. (2024)) by a stoma with distinct dorsal tooth (*vs* small hook-shaped dorsal tooth), smaller ratio of body length to neck length (b) (4.8-5.7 vs. 5.8-6.6), smaller ratio of body length to tail length (c) (13.6-15.0 vs. 20.2-22.8), as well as stoma (8.5-8.9 µm vs. 11.3-12.9 µm) and isthmus length (34.1-41.3 µm vs. 26.9-30.9 µm).

From *P. kolymaensis* (after Shatilovich et al. (2023)) specifically by gonochoristic reproduction mode indicated by the abundance of males.

From *P. subelongatus* (after Mehdizadeh et al. (2013)) by a differently shaped lip region with all lips equal in size (*vs* lips completely amalgamated in pairs), stoma with distinct dorsal tooth (*vs* without dorsal tooth), larger ratio of body length to neck length (b) (4.8-5.7 vs. 4.1-4.5) and a smaller ratio of body length to tail length (c) (13.6-15.0 vs. 19.4-21.6), a wider body width (mid: 43.1-47.9 µm vs. 27.0-36.6 µm; anus: 26.2-31.0 µm vs. 18.0-21.6 µm), as well as a shorter stoma length (8.5-8.9 µm vs. 10.8-13.0 µm) and a shorter distance of excretory pore to anterior end (119.1-133.8 µm vs. 137.7-144.5 µm).

From *P. rigidus* (after Seddiqi et al. (2016)) by a stoma with distinct dorsal tooth (*vs* without dorsal tooth), smaller ratio of body length to body diameter (a) (17.4-20.3 vs. 24.3-27.1), a smaller ratio of body length to neck length (b) (4.8-5.7 vs. 6.0-6.4) and a smaller ratio of tail length to tail diameter (c’) (2.0-2.2 vs. 3.1-3.7), a larger body width (mid: 43.1-47.9 µm vs. 30.2-34.4 µm; anus: 26.2-31.0 µm vs. 16.4-18.6 µm), as well as a longer corpus length (86.6-99.0 µm vs. 73.5-78.3) and a shorter spicule length in males (26.5-30.1 µm vs. 30.6-32.6 µm).

From *P. orientalis* (after Seddiqi et al. (2016)) by a stoma with distinct dorsal tooth (*vs* with small dorsal tooth), phasmid located at posterior 1/3rd of tail length in females (*vs* at middle of tail), larger ratio of body length to tail length (c) (13.6-15.0 vs. 10.7-12.3), smaller ratio of tail length to tail diameter (c’) (2.0-2.2 vs. 2.9-3.5) and broader body width (mid: 43.1-47.9 µm vs. 29.2-40.2 µm; anus: 26.2-31.0 µm vs. 14.9-19.5 µm), as well as longer corpus length (86.6-99.0 µm vs. 71.9-78.3 µm). Original description of *Panagrolaimus orientalis* (Korentchenko, 1986) specifies an anisomorphic stoma with two subdorsal teeth – a condition that is different from both *P. nebliphilus* sp. nov. and Seddigi’s redescription.

From *P. facetus* (after Mehdizadeh et al. (2013)) by a stoma with distinct dorsal tooth (*vs* without dorsal tooth), phasmid located at posterior 1/3rd of tail length in females (*vs* at anterior 1/3rd of tail), smaller ratio of body length to body diameter (a) (17.4-20.3 vs. 23.2-29.8), ratio of body length to tail length (c) (13.6-15.0 vs. 12.5-13.5), ratio of tail length to tail diameter (c’) (2.0-2.2 vs. 2.3-3.7) and body width (mid: 43.1-47.9 µm vs. 25.1-33.3 µm; anus: 26.2-31.0 µm vs. 15.1-25.7 µm), as well as the stoma length (8.5-8.9 µm vs. 12.1-13.1 µm).

From *P. labiatus* (after Yadav et al. (2022)) by a stoma with distinct dorsal tooth (*vs* without dorsal tooth), phasmid located at posterior 1/3rd of tail length in females (*vs* at anterior 1/3rd of tail), smaller ratio of body length to body diameter (a) (17.4-20.3 vs. 20.4-23.6), larger ratio of body length to tail length (c) (13.6-15.0 vs. 11.2-13.4) and smaller ratio of tail length to tail diameter (c’) (2.0-2.24 vs. 2.9-3.7), a broader body width (mid: 43.1-47.9 µm vs. 32.0-40.0 µm; anus: 26.2-31.0 µm vs. 17.7-24.3 µm), as well as a longer corpus (86.6-99.0 µm vs. 73.9-86.1 µm).

From *P. concolor* (after Mehdizadeh et al. (2013)) by a stoma with distinct dorsal tooth (*vs* with small dorsal tooth), narrower body width mid-body (43.1-47.9 µm vs. 49.0-52.6 µm), a longer corpus (86.6-99.0 vs. 78.4-84.8 µm) and a shorter distance of excretory pore to anterior end (119.1-133.8 µm vs. 149.2-161.8 µm).

From *P. papillosus* (after Mehdizadeh et al. (2013)) by a smaller ratio of body length to body diameter (a) (17.4-20.3 vs. 22.7-24.1), a smaller ratio of body length to tail length (c) (13.6-15.0 vs. 19.2-20.4) and a larger ratio of tail length to tail diameter (c’) (2.0-2.2 vs. 1.7-1.9), as well as a broader body width (mid: 43.1-47.9 µm vs. 31.2-34.8 µm; anus: 26.2-31.0 µm vs. 20.0-22.8 µm) and shorter stoma length (8.5-8.9 µm vs. 11.4-13.2 µm). Type population of *P. papillosus* described by Loof is most likely asexual (Loof, 1971) while both *P. nebliphilus* and Mehdizadeh’s population are gonochoristic

From *P. leperisini* (after Massey (1974); Girgan et al. (2018)) by a papilliform anterior sensilla (*vs* setiform), smaller ratio of body length to body diameter (a) (17.4-20.3 vs. 29.5-34.9), a smaller ratio of body length to tail length (c) (13.6-15.0 vs. 18.9-19.2) and a smaller ratio of tail length to tail diameter (c’) (2.0-2.2 vs. 2.5-2.6), as well as a broader body width (mid: 43.1-47.9 µm vs. 31.0-33.0 µm; anus: 26.2-31.0 µm vs. 18.0-21.0 µm) and shorter corpus length (86.6-99.0 µm vs. 107.0-114.0 µm).

### Genome assemblies and phylogeny

#### Genome assemblies

All three species identification are supported by high-quality genomes assemblies. The *P. einhardi* sp. nov. genome was further refined using Hi-C technology. The chromosome-level genome assembly of *P. einhardi* sp. nov. includes 44 scaffolds, with a N50 of 28 Mb and an N90 of 24 Mb (Table 2). 99.86 % of the assembly is in scaffolds longer than 50 kb. The total genome size is 116 Mb, with a GC content of 28.38 %. According to the BUSCO Nematoda odb12 lineage, 96.3 % of orthologs are complete, with 5.5 % duplicated, 0.8 % fragmented, and 2.9 % missing (Figure 8A, Supplementary Material Figure S21, S24).

The following two assemblies were generated using Nanopore sequencing. The genome of *P. shuimeiren* sp. nov. is comparatively smaller, totaling 69 Mb, with a N50 of 13 Mb and an N90 of 12 Mb, consisting of 49 scaffolds. 99.19 % of the assembly is in scaffolds longer than 50 kb. Its GC content is 36.48 %, with 97.8 % BUSCO completeness and a duplication rate of 7.2 %. Fragmented orthologs account for 0.7 %, and 1.5 % are missing (Figure 8B, Supplementary Material Figure S22, S25).

**Figure 8.**
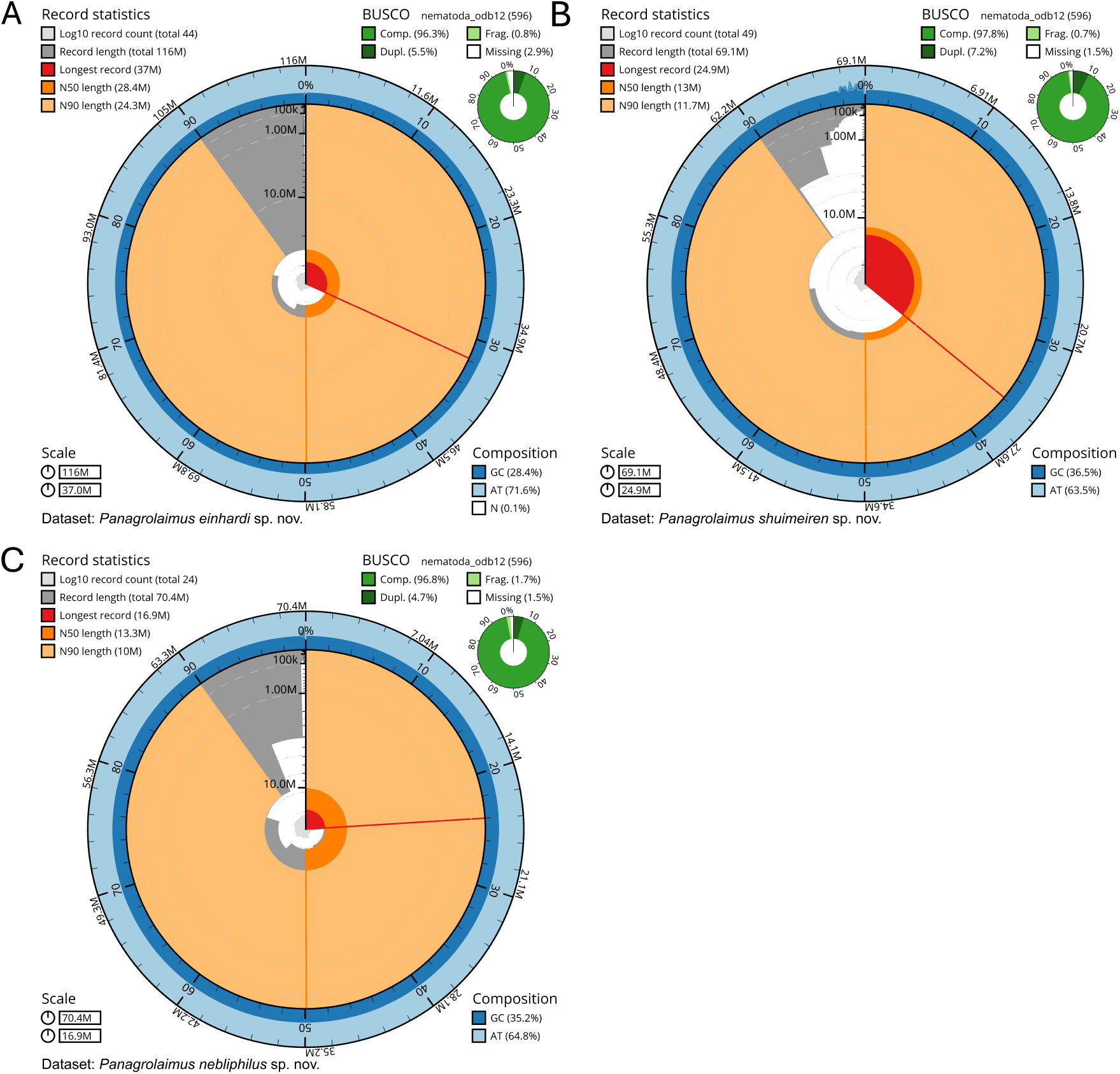
Snail plots summarize assembly statistics from BlobToolKit. The longest scaffold is marked with a red line, and the N50 value is highlighted by an orange line, while the N90 value is shown as a light orange pie chart. The outer circle illustrates GC and AT composition in dark and light blue, respectively. BUSCO analysis using the insecta odb12 database provides details on completeness, fragmentation, duplication, and missing genes in the upper right corner. Blobplots and k-mer spectra in Supplementary Material Figure S21-S26. A, *Panagrolaimus einhardi* sp. nov. B, *Panagrolaimus shuimeiren* sp. nov. C, *Panagrolaimus nebliphilus* sp. nov.

*P. nebliphilus* sp. nov. has similar assembly size compared to *P. shuimeiren* sp. nov. with approximately 70 Mb, with 24 scaffolds, a N50 of 13 Mb, and an N90 of 10 Mb. 99.82 % of the assembly is in scaffolds longer than 50 kb. Its GC content is 35.19 %. BUSCO scores indicate 96.8 % complete genes, with a duplication rate of 4.7 %. Fragmentation affects 1.7 %, and 1.5 % are missing (Figure 8C, Supplementary Material Figure S23, S26). Repeat content of the three assemblies is highest for *P. einhardi* sp. nov. with 13.6 % and lowest for *P. nebliphilus* sp. nov. with 13.1 % (Table 2).

**Table 2.** Summary for each assembly, indicating total size, N50, N90, GC and repeat content, the longest scaffold, and the number of scaffolds.

| Assembly | *P. einhardi* | *P. shuimeiren* | *P. nebliphilus* |
| --- | --- | --- | --- |
| Assembly size (bp) | 116,281,119 | 69,123,706 | 70,365,442 |
| N50 (bp) | 28,424,827 | 13,020,021 | 13,308,294 |
| N90 (bp) | 24,316,838 | 11,656,429 | 10,040,455 |
| GC content (%) | 28.38 | 36.48 | 35.19 |
| Repeats content (%) | 13.6 | 13.2 | 13.1 |
| BUSCO<br>(nematoda_odb12) | C:96.3%[S:90.8% D:5.5%]<br>F:0.8% M:2.9%<br>n:596 E:6.6% | C:97.8%[S:90.6% D:7.2%]<br>F:0.7% M:1.5%<br>n:596 E:6.3% | C:96.8%[S:92.1% D:4.7%]<br>F:1.7% M:1.5%<br>n:596 E:6.1% |
| Longest scaffold (bp) | 37,040,167 | 24,873,950 | 16,917,494 |
| No. of scaffolds | 44 | 49 | 24 |

Pairwise whole-genome alignments between the three assemblies showed the contiguity of *P. einhardi*’ sp. nov. assembly (Figure 9B, Supplementary Material Figure S27). It is generally expected that diploid *Panagrolaimus* species have four chromosome pairs. Chromosomes have been labeled according to conserved chromosome elements, called Nigon elements (Tandonnet et al., 2019), as assigned by Guiglielmoni et al. (in preparation). The contiguity of *P. einhardi* sp. nov. genome is also displayed by the length of the longest scaffold, where 37 Mb is approximately 32 % of the full assembly. For *P. shuimeiren* sp. nov. the longest scaffold with 25 Mb represents 36 % of the assembly and for *P. nebliphilus* sp. nov. the longest scaffold represents 24 % (16.9 Mb) of the assembly.

**Figure 9.**
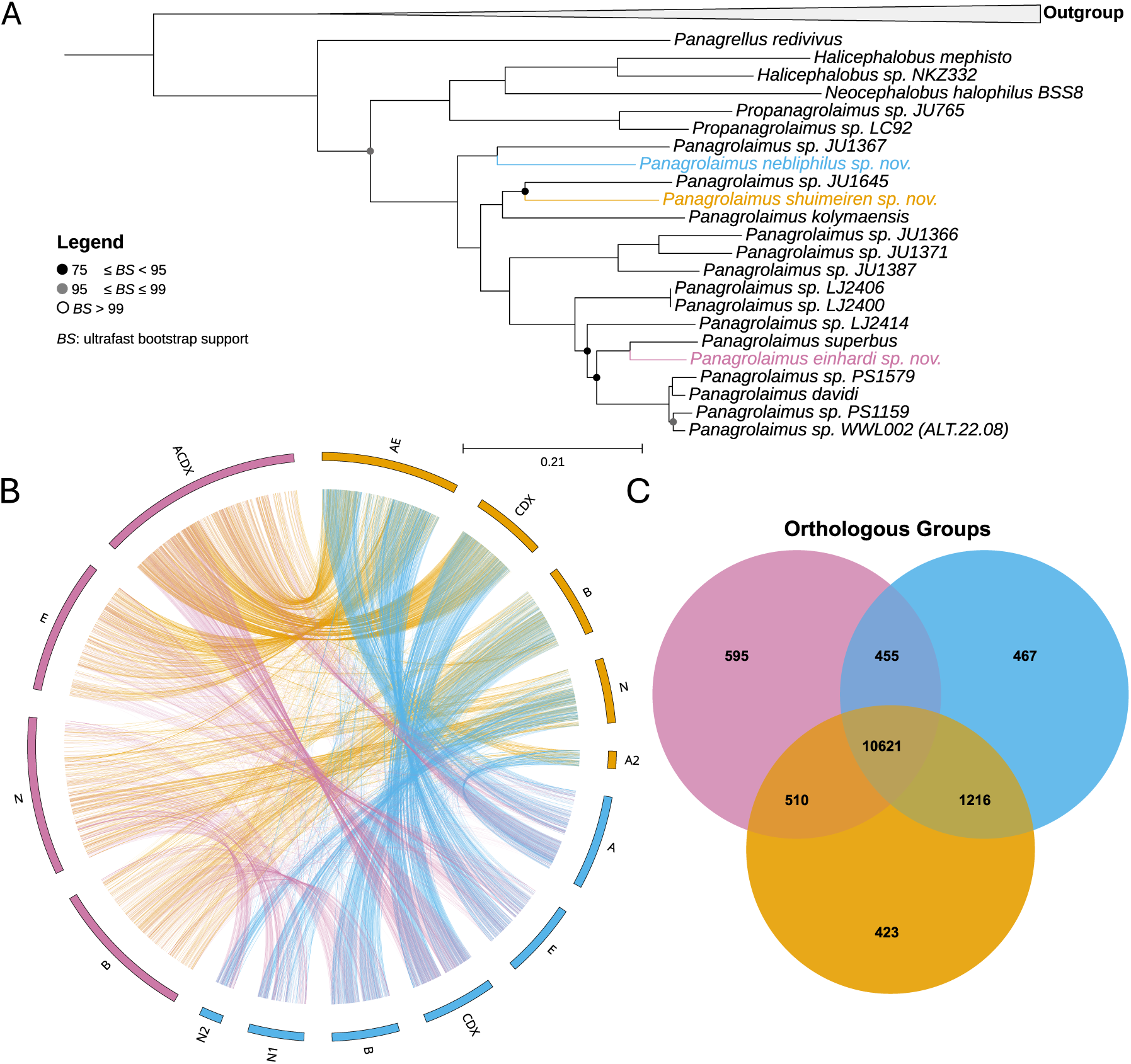
A, Phylogenetic tree using maximum-likelihood (ML) based on ultra-conserved elements (UCEs) with 1,000 ultrafast bootstraps. Bootstrap values between 75 to 95 are indicated with black circles, bootstrap values between 95 and 99 are indicated with grey circles. No circle refers to a bootstrap value over 99. Low supported bootstrap values (lower than 75) were collapsed. The outgroup was used to root the tree and consists of four *Acrobeloides* species: *A. obliquus*, *A. thornei*, *A. tricornis* PAP.22.17 (WWL709) and *A. maximus*. B, Circos plot visualizing pairwise alignments between the assemblies. Smaller scaffolds are not displayed. Scaffold names follow the Nigon element assignment proposed by Guiglielmoni et al. (in preparation). C, Venn diagram of orthologous protein clusters. The distribution of orthologous protein clusters among the analyzed species is visualized, with each circle representing a species. Overlapping regions indicate gene families shared between two or more species. The numbers indicate the size of each cluster set. Created with OrthoVenn3. The colors indicate the species in all three images: Pink = *P. einhardi* sp. nov., orange = *P. shuimeiren* sp. nov., blue = *P. nebliphilus* sp. nov.

A comparative analysis of the three genomes was conducted using OrthoVenn3, which identified both shared and unique orthologous clusters. The distribution of these clusters across the genomes is illustrated in the Venn diagram (Figure 9B, Supplementary Material Figure S28). A total of 58,694 protein sequences were analyzed, resulting in the identification of 14,287 orthologous clusters, of which 7,565 were single-copy clusters. Among the three genomes, 10,621 clusters (74.34 % of all clusters) were identified as shared (Figure 9C), indicating a conserved core set of genes. Pairwise comparisons revealed distinct relationships: there were 1,216 clusters (8.51 %) shared between *P. nebliphilus* sp. nov. and *P. shuimeiren* sp. nov., 510 clusters (3.57 %) shared between *P. shuimeiren* sp. nov. and *P. einhardi* sp. nov., and 455 clusters (3.18%) shared between *P. nebliphilus* sp. nov. and *P. einhardi* sp. nov. (Figure 9C). These findings reflect differences in evolutionary relationships, indicating that *P. nebliphilus* sp. nov. and *P. shuimeiren* sp. nov. share the most orthologous clusters, while *P. nebliphilus* sp. nov. and *P. einhardi* sp. nov. share the fewest. OrthoVenn3 not only identified shared clusters among the genomes but also revealed unique clusters specific to individual species. Specifically, there were 595 clusters (4.16 %) unique to *P. einhardi* sp. nov., 467 clusters (3.27 %) unique to *P. nebliphilus* sp. nov., and 423 clusters (2.96 %) unique to *P. shuimeiren* sp. nov. (Figure 9C). The analysis identified the presence of singletons, which accounted for an overall percentage of 5.35 %. This corresponds to a total of 3,140 singletons found across the examined sequences. Specifically, in the case of *P. einhardi* sp. nov., there were 1,112 singletons out of 20,594 protein sequences. For *P. shuimeiren* sp. nov., 1,092 singletons were detected from a total of 19,460 protein sequences. Lastly, *P. nebliphilus* sp. nov. had 936 singletons identified out of 18,640 protein sequences.

#### Ultra conserved elements based phylogeny

Harvesting of UCEs from the three new assemblies led to 1488 elements for *P. einhardi* sp. nov., 1391 elements for *P. shuimeiren* sp. nov. and 1402 elements for *P. nebliphilus* sp. nov. from a total of 1606 UCEs. An alignment of the UCEs was created including 27 taxa. Of the alignment, 14.5 % were identified as invariable sites. A phylogenetic tree using Maximum-Likelihood with 1000 ultrafast bootstraps (Figure 9A) was created with four *Acrobeloides* species as an outgroup clade (accession numbers in Supplementary Material Table S1). The analysis revealed that the closest relatives of *P. einhardi* sp. nov. are *P. superbus* and *P. davidi*, with good bootstrap support for the shared node at 93 % and for the preceding node at 87 %. *P. shuimeiren* sp. nov. and *P. nebliphilus* sp. nov. were found to be relatively closely related. *P. shuimeiren* sp. nov. formed a clade with strain JU1645, showing 98 % support, and positioned adjacent to *P. kolymaensis*, which had very strong support (100 %). In contrast, *P. nebliphilus* sp. nov. was resolved more basally than *P. kolymaensis*, along with strain JU1367. The complete *Panagrolaimus* clade, consisting of 17 members, received maximal support (100 %).

From the assemblies, 18S rRNA sequences were extracted and consensus sequences were created. Based on traditional methods, a phylogenetic tree using this single locus was reconstructed building upon the analysis of Abolafia (2025) (Figure 10). Instead of Bayesian inference, we here used Maximum likelihood to match the analysis of UCEs. The entire clade encompassing *Microlaimus* and *Panagrolaimus* is supported at 93 %. Notably, “Clade I”, which incorporates *Macrolaimus* species, is not resolved as a single clade compared to Abolafia (2025), but rather is subdivided, with *Macrolaimus ruehmi* situated more basally relative to *M. crucis* and *M. canadensis*. Apart from that, the topology is similarly resolved with the addition of new 18S sequences derived from genome assemblies incorporated into the phylogenetic tree. The placements of *Panagrellus redivius*, *Turbatrix aceti*, *Neocephalobus* species, and *Propanagrolaimus* species are supported with maximal confidence. The *Halicephalobus* clade has 92 % support, and the distinction between *H. gingivalis* and *Halicephalobus* sp. NKZ332 is supported at 76 %. The 18S phylogeny (Figure 10) places the species described here at similar positions as the UCE phylogeny (Figure 9A). *P. einhardi* sp. nov. is reaffirmed within the same clade as *P. superbus* (Clade II, 92 %). *P. nebliphilus* sp. nov. is also positioned alongside *Panagrolaimus* sp. JU1367 within “Clade III,” which additionally includes *P. rigidus* and *P. labiatus*. The bifurcation between “Clade III” and “Clade IV” has support at 90 %. *P. shuimeiren* sp. nov. is again associated with *Panagrolaimus* sp. JU1645 (77 %) and exhibits the closest relationship to *P. kolymaensis* (97 %). Overall, the bootstrap support values for the 18S rRNA-based phylogenetic tree (Figure 10) are comparatively lower and less robust than those observed in the UCE-based phylogeny (Figure 9A), with most of the dichotomies in the uncollapssed tree poorly supported.

**Figure 10.**
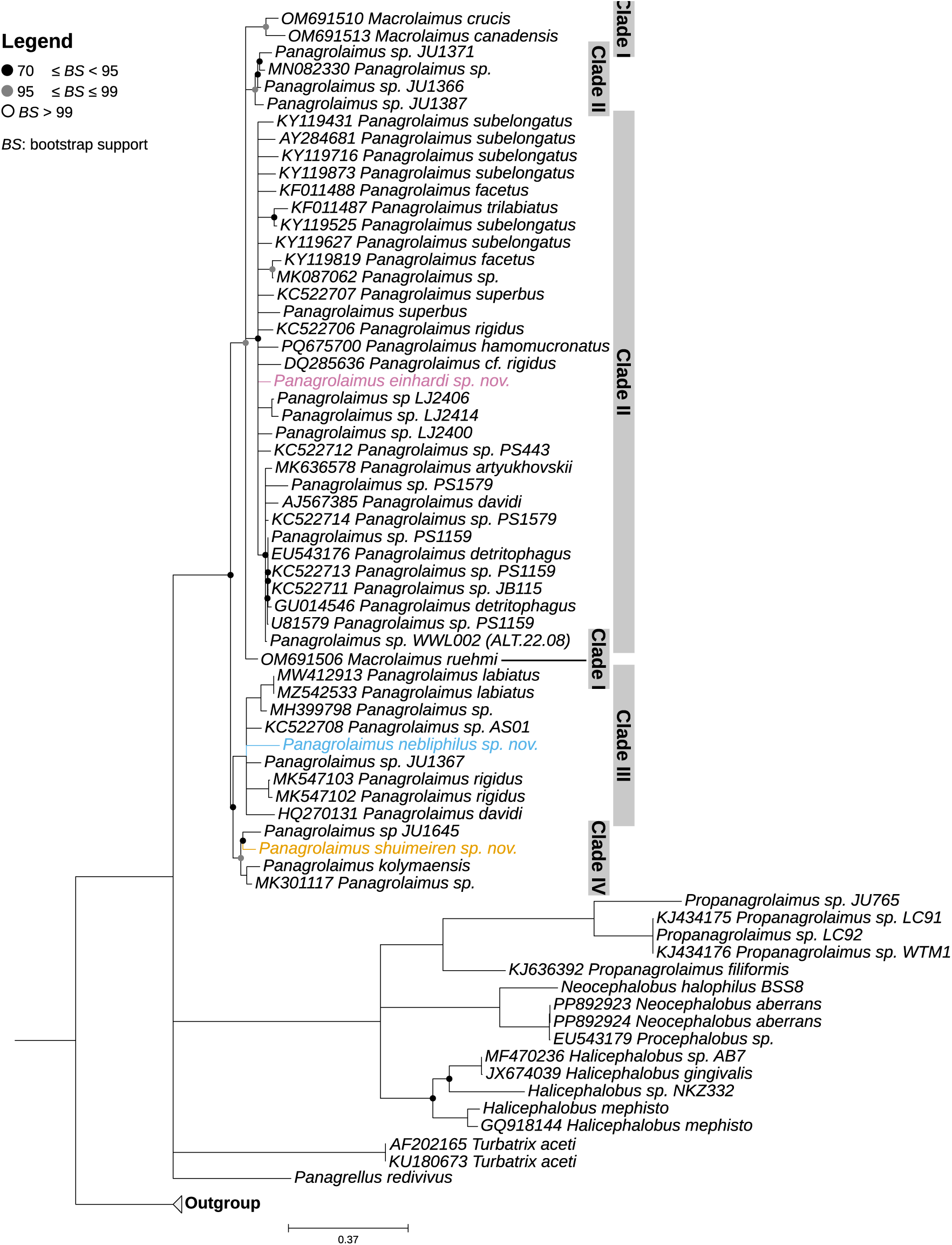
Phylogenetic tree using maximum-likelihood (ML) inference based on 18S rRNA with 1,000 bootstraps replicates based on Abolafia (2025) with adddition of new sequences. Bootstrap values between 70 to 95 are indicated with black circles, bootstrap values between 95 and 99 are indicated with grey circles. No circle refers to a bootstrap value over 99. Clades with low support (bootstrap values lower than 70) were collapsed. The outgroup was used to root the tree and consists of four *Acrobeloides* species: *A. obliquus*, *A. thornei*, *A. tricornis* PAP.22.17 (WWL709) and *A. maximus*. The colors indicate the species in all three images: Pink = *P. einhardi* sp. nov., orange = *P. shuimeiren* sp. nov., blue = *P. nebliphilus* sp. nov.

## DISCUSSION

The family Panagrolaimidae is ecologically a highly diverse family of nematodes sharing a uniform body plan, hindering accurate morphological species identification. Describing species within the genus *Panagrolaimus* is particularly challenging, as species descriptions rely on fine details that can be difficult to obtain (Abolafia and Peña-Santiago, 2006; Abolafia, 2025) and eventually carry sampling biases. Accurate morphological measurements are essential for proper species description, but not necessarily decisive. In this work, we present the unique morphology of three species that can be distinguished from others within the genus. The complementary use of morphology and genomics provides a solid framework for refining the taxonomic resolution of the described species. It highlights the value of expanding genomic resources for future systematic research.

Among the three species, the most significant morphological differences were observed in the body width at mid-body and the anus, which clearly distinguished the species from one another. *P. shuimeiren* sp. nov. has the narrowest body, while *P. nebliphilus* sp. nov. features the widest body. Additionally, *P. shuimeiren* sp. nov. has an overall shorter body length compared to the other two species, a difference that is also noticeable in the length of the pharynx and the distance from the excretory pore to the anterior end. Furthermore, *P. nebliphlus* sp. nov. and *P. shuimeiren* sp. nov. differ in the ratio of body length to the greatest body diameter (a). However, it must be remembered that all measurements can be subject to errors and can be affected by fixation artifacts (shrinkage in higher than optimal concentration of formaldehyde-based fixatives, or when dehydration to absolute glycerol happens too fast) or nematodes being squashed by a cover slip.

Differences in the genital papillae (GP) have also been demonstrated to be among the most reliable diagnostic features with six GP in males of *P. einhardi* sp. nov. and six GP and one midventral papillae (MP) in *P. shuimeiren* sp. nov. Male genital papillae patterns showed notable divergence in *P. nebliphilus* sp. nov. with eight GP and one MP.

Phylogenetically, the closest relative of *P. einhardi* sp. nov. is *P. superbus*; however, they exhibit several distinct differences in their morphological and morphometric traits. *P. einhardi* sp. nov. has a narrower body width (at mid-body and anus) and shows pronounced differences in the ratio of body length to tail length (c), as well as the ratio of tail length to tail diameter at the anus (c’). Additionally, there are differences in the distance from the excretory pore to the anus and in the length of the stoma. In males, *P. einhardi* sp. nov. has a shorter gubernaculum and spicule lengths, along with two pre-cloacal genital papillae, compared to three in *P. superbus*. The lip region is also different between two species. Furthermore, *P. einhardi* sp. nov. differs from the related species *P. davidi* by having a shorter basal bulb length and two pre-cloacal pairs of genital papillae in males, whereas *P. davidi* has only one, in addition to the latter species being endemic to Antarctica. *P. nebliphlus* sp. nov. and *P. shuimeiren* sp. nov. are distinct on the phylogenetic tree, with one of their closest relatives being recently discovered and described *P. kolymaensis*. However, they are clearly differentiated from *P. kolymaensis* based on their reproductive modes, as both are gonochoristic compared to the parthenogenetic *P. kolymaensis*.

The morphological differences still suggest a considerable degree of similarity within the genus *Panagrolaimus* and it is crucial to acknowledge that certain specimens may display phenotypic variability stemming from uncertainties in environmental or experimental conditions, as noted by Yadav et al. (2022); Mianowska (1977) in the case of *Panagrolaimus*. Moreover, many redescriptions of known species do not match the original descriptions in certain morphological or biological features. For example, *P. papillosus* described by Mehdizadeh et al. (2013) is gonohoristic while *P. papillosus* as originally described by Loof (1971) is asexual. Similarly, redescriptions of *P. orientalis* by Seddiqi et al. (2016) differs from original specimens of Korentchenko (1986) in stoma morphology. *P. labiatus* described by Mehdizadeh et al. (2013) and *P. labiatus* described by Yadav et al. (2022) are also not the same species, differing in multiple characters related to female and male tail at least, position of phasmid and shape of spicules. Population of *P. subelongatus* described by Mehdizadeh et al. (2013) has no teeth in the gymnostom, while the population described by Rühm (1956) has small teeth on all three rhabdia, the arrangement of papilla on male tail is different, and the population described by Mehdizadeh et al. (2013) has longer spikes. Populations of *P. superbus* described by Abolafia and Peña-Santiago (2006) and Mehdizadeh et al. (2013) differ in the shape of the lip region and the position of the phasmid in females. These examples highlight the necessity for rigorous re-evaluation of existing species descriptions of various *Panagrolaimus* species, re-evaluation of remaining type specimens, re-collection of specimens from type localities and described vouchers, and additional taxonomically useful features (morphological, metric, and genomic) to improve species descriptions within this genus.

Sequencing of short DNA sequences, such as 18S rDNA and 28S rDNA, has been widely utilized in nematodes (Eyualem and Blaxter, 2003; Shatilovich et al., 2023; Rawson et al., 2024; Blaxter, 2011) and has provided some evidence of hidden genetic diversity between morphologically indistinguishable populations (Figure 10). However, difficulties in DNA barcoding of nematodes have been identified, including misidentified specimens in databases, and differing genetic markers that cause inconsistencies in phylogenetic placements (Eyualem and Blaxter, 2003; Rawson et al., 2024; Villegas et al., 2025). One example of misclassifications is *Propanagrolaimus* sp. JU765, which has historically been assigned to the genus *Panagrolaimus* and is evidenced by 18S and 28S ribosomal RNA gene sequences available in NCBI GenBank database, which still reflect the outdated taxonomy. Some of these challenges have been addressed by the establishment of a curated database for 18S sequences (Gattoni et al., 2023), but experience shows that such databases are rarely maintained for a long period of time. Although 18S rRNA remains a standard marker for phylogeny across the entire phylum Nematoda as well as for barcode-based identification, our results show that its resolution within the genus *Panagrolaimus* is limited compared to genome-derived UCEs. The 18S-based phylogeny aligned closely with the UCE phylogeny in classifying the newly described species; however, numerous relationships had weak bootstrap support or were inconsistently resolved, compared to the phylogeny based on UCEs. In particular, the *Macrolaimus* clade was not found as a monophyletic clade, unlike in Abolafia (2025). The UCE phylogeny consistently provided higher support values and a clearer resolution of interspecific relationships within *Panagrolaimus*. At the same time, the phylogeny based on UCEs is more balanced in terms of difference in substitution rates expressed as individual branch length, between *Panagrolaimus* and its close relatives, compared to the 18Ss-based phylogeny. Both emphasize the benefit of using hundreds of independent genomic loci instead of a single conserved marker, which can also be distorted by intragenomic polymorphism (Wang et al., 2023). While 18S rRNA sequences extracted from genome assemblies confirm phylogenetic placement of newly discovered species, UCEs provide significantly greater resolution and reliability for distinguishing species and making evolutionary inferences in *Panagrolaimus*, as also supported by Villegas et al. (2025). This is also further backed by the similar studies conducted in *Caenorhabditis* phylogeny (Slos et al., 2017; Stevens et al., 2019). Compared to short DNA sequences, genomes contain a mix of both conserved and variable regions, and they also encode extensive information about a species’ development and evolutionary history, which can support various analytical methods related to genetic functions and traits, making it beneficial to publish species descriptions alongside genome assemblies (Stevens et al., 2019), even if these assemblies are partial or fragmented.

In this study, we obtained high-quality assemblies, supported by high BUSCO scores. The recently released twelfth version of the database (nematoda odb12) (Tegenfeldt et al., 2024) now includes 54 nematode species, a considerable increase from the seven species featured in the previous version (nematoda odb10). This significantly enhances the reliability of detected orthologs, with over 96 % completeness observed in all three assemblies. Orthologs refer to genes present in different species that have descended from a common ancestral gene, usually maintaining similar functions (Sun et al., 2023; Wang et al., 2015). While newly described species share characteristic morphological structures of the genus, genetic analysis reveals unique ortholog clusters and well-supported phylogenetic branches, reflecting their relatively distant relationships and possibly diverging biology. A core group of 10,621 clusters shared by all three species was detected with OrthoVenn3 (Sun et al., 2023), indicating highly conserved genes likely involved in crucial biological processes. Additionally, each species retained unique clusters. These species-specific clusters may represent expansions of gene families or functions that are unique to lineages due to ecological or physiological adaptations (Wang et al., 2015). The existence of distinct clusters further emphasizes the need to classify these organisms as separate species, thereby highlighting their unique traits and ecological roles within their respective environments. Although they share a conserved genetic core, each lineage has developed distinct genes and functions due to evolutionary divergence, adaptation to specific environments, or the development of particular traits. Comparisons between pairs of species also indicate that 510 clusters are shared by *P. einhardi* sp. nov. and *P. shuimeiren* sp. nov., 455 clusters are shared by *P. einhardi* sp. nov. and *P. nebliphilus* sp. nov., and 1,216 clusters are shared by *P. shuimeiren* sp. nov. and *P. nebliphilus* sp. nov. The varying sizes of these overlaps suggest different levels of evolutionary relatedness, with larger overlaps pointing to a more recent common ancestry. Notably, both *P. shuimeiren* sp. nov. and *P. nebliphilus* sp. nov. inhabit desert habitats, indicating they may have needed similar ecological or physiological adaptations. In conclusion, we identified both conserved core genes that link these species, but also divergent clusters that indicate the genetic basis of their ecological and evolutionary distinctions.

In the phylogenetic tree, both *P. nebliphilus* sp. nov. and *P. shuimeiren* sp. nov. are closely related and belong to a group of more basal *Panagrolaimus*. Their proximity to the ancient *P. kolymaensis* suggests that these two species are relatively old, likely due to the stable conditions in their desert habitats, suggesting that both species have survived for prolonged times in extreme conditions. Notably, we find related species in very different habitats that differ in geographic location, as well as climatic conditions, but all appear to be “dry” either due to cold (cryobiotic) or hot (anhydrobiotic) coonditions. *P. kolymaensis* was retrieved from the deeply frozen layers of Russian permafrost (Shatilovich et al., 2023), highlighting its adaptation to extremely cold environments, with water supply being limited by freezing. In contrast, *Panagrolaimus* sp. JU1645 originated from Cape Verde, where it thrives in a semi-arid climate characterized by limited rainfall and higher temperatures (McGill et al., 2015). Similarly, *Panagrolaimus* sp. JU1367 is found in Tamil Nadu, India, specifically in regions characterized by a dry sub-humid to sub-arid climate, with distinct dry and wet seasons (McGill et al., 2015). The newly described *P. nebliphilus* sp. nov. originates from the Atacama Desert, together with *P. shuimeiren* sp. nov. from the Namib Desert, two of the known driest areas on Earth, with regular but very limited water supply (Hartley et al., 2005; Huntley, 2023). *P. nebliphilus* sp. nov. and *P. shuimeiren* sp. nov. reproduce sexually, a characteristic also observed in *Panagrolaimus* sp. JU1645 and *Panagrolaimus* sp. JU1367 (McGill et al., 2015). In contrast, *P. kolymaensis* reproduces parthenogenetically. Since other asexual *Panagrolaimus* species appear later in the phylogenetic tree, the loss of sexual reproduction in *P. kolymaensis* may have resulted from an independent evolutionary event and may not be related to the environmental conditions that this species is assumed to have lived in naturally. For more recent parthenogenetic species, a hybridization event is suspected to be the origin, a hypothesis that also applies to *P. kolymaensis* (Shatilovich et al., 2023).

*P. einhardi* sp. nov. is characterized as a relatively younger species in the phylogenetic tree when compared to the other two newly described species. *P. einhardi* sp. nov. shows a close genetic relationship with *P. superbus*, which is known to inhabit the Arctic regions (Boström, 1988), and *P. davidi* (Wharton, 1998), found in the Antarctic. In contrast to these related species, which thrive in extremely cold environments, *P. einhardi* sp. nov. is found in the temperate climates of Germany.

Collectively, all of these species demonstrate a remarkable diversity of ecological niches and adaptability of *Panagrolaimus*. The adaptability to various environmental conditions are providing important insights into the evolutionary trajectories that facilitate survival in diverse habitats while maintaining morphological traits. This adaptability may reveal how certain species navigate ecological challenges, preserving their structural characteristics through evolutionary pressures.

Morphological analysis remains essential in taxonomy and evolutionary studies, offering direct insights into observable and quantifiable phenotypic variation. Nonetheless, morphology can be confounded by convergent evolution, the presence of cryptic species, or phenotypic plasticity, which may complicate accurate classification (Schneider and Meyer, 2016; Losos, 2011). Particularly since it is assumed that the initial phases of speciation primarily influence physiological, immunological, reproductive, or behavioral characteristics more than morphological traits (Struck et al., 2018). Thus, both comparative genomic studies and taxonomic investigation of closely related species, require high-quality reference genomes, especially when aiming to understand complex life traits that may be caused by subtle genomic differences, which is particularly valuable for functional studies (Hellekes et al., 2023).

Genomics enable detailed mapping of genetic variation across individuals and taxa, which is powerful for resolving deeper evolutionary relationships, detecting hidden diversity, gene flow or hybridization. Increasing the number of genome assemblies has potential to increase taxonomic resolution in cryptic genera like *Panagrolaimus* and help name unclassified laboratory strains. However, producing high-quality assemblies can be difficult due to challenges of working with non-model nematodes, such as obtaining sufficient, low-fragmentation DNA. Pooling multiple individuals is often necessary, but heterozygosity and bacterial contamination can complicate the assembly process (Guiglielmoni et al., 2022). Advances in DNA extraction protocols and long-read sequencing now improve genome assemblies of non-model invertebrates, like *Panagrolaimus* and *Romanomermis* (Guiglielmoni et al., 2024). Other challenges here include high costs, significant technical requirements, and complex bioinformatic analyses, as well as gaps in understanding the links between genotype and ecologically relevant phenotypes (Rhie et al., 2021; Guiglielmoni et al., 2022; Lehner, 2013).

An integrative approach combining both morphological and genomic data in taxonomy provides enhanced resolution in species delimitation and evolutionary inference. It is important to note, that not every described species currently possesses a genome assembly, limiting this approach. Initiatives such as the Earth Biogenome Project (Gupta, 2022) aim to generate high-quality reference genome sequences for all 1.8 million eukaryotic species, but achieving this goal still appears to be unrealistic for many working with species and hyperdiverse groups of organisms, like nematodes. Starting small, focusing on a single lineage, and using it to develop reliable and affordable tools and protocols for generating high-quality genomes is a viable alternative, as the results of this project demonstrate. Building upon and expanding, instead of a “grab and go” approach of phylum/kingdom/everything-wide initiatives may be a more realistic and reliable way forward in the long run.

Genomes encapsulate the vital information needed to create an individual organism. By weaving together morphology and genomics, we can unlock a deeper understanding of complex evolutionary histories, ultimately illuminating the rich biological diversity of morphologically uniform and seemingly unremarkable for a “classic taxonomist” cryptic families, such as the Panagrolaimidae. Morphological traits anchor genomic diversity in observable phenotypes, but so are the physiological and biochemical traits that are not visible under the optical or electron microscope, but nonetheless play an undeniable role in species survival. They all identify what a species is. Thus, comparative genomic analysis is actually the simplest currently available tool that can clarify relationships where morphological data alone are ambiguous or confounded.

## CONCLUSION

*Panagrolaimus* comprises numerous cryptic species that exhibit significant morphological similarity yet possess considerable ecological, genetic, and genomic diversity. This is particularly evident in species inhabiting extreme environments, such as the Atacama and Namib Desert. In this publication, we describe three novel species: *P. einhardi* sp. nov., *P. nebliphilus* sp. nov., and *P. shuimeiren* sp. nov. *P. einhardi* sp. nov. has frequently been referred to as *Panagrolaimus* sp. ES5 in various research studies. The predominance of strain numbers over formal species names in *Panagrolaimus*, just like in some other intensively studied nematode lineages, highlights the necessity for enhanced and more precise taxonomic classification within this genus.

## Supporting information

Supplementary Results

## ACKNOWLEDGMENTS

We thank the ITCC (IT Center University of Cologne) for providing compute resources on the DFG-funded HPC (High Performance Computing) system RAMSES (Research Accelerator for Modeling and Simulation with Enhanced Security) as well as support (DFG funding number: INST 216/512-1 FUGG). We would like to thank the Wellcome Sanger Institute, the Cologne Center for Genomics (CCG), and the Genomics and Transcriptomics Laboratory (GTL) for their support.

## AUTHOR CONTRIBUTIONS

Laura C. Pettrich (Conceptualization, Methodology, Investigation, Data curation, Formal analysis, Writing – original draft), Arunee Suwanngam (Investigation, Data curation, Formal analysis, Writing – original draft), Nadège Guiglielmoni (Methodology, Investigation, Data curation, Formal analysis), Laura I. Villegas (Methodology, Investigation), Amy Treonis (Methodology, Investigation, Data curation), Noelle Ledoux (Methodology, Investigation, Data curation), Lewis Stevens (Methodology, Investigation), Manuela Kieninger (Methodology, Investigation), Michael Paulini (Data curation), Mark Blaxter (Supervision, Funding acquisition), Ann-Marie Waldvogel (Conceptualization, Writing – review & editing, Supervision, Project administration, Funding acquisition), Oleksandr Holovachov (Conceptualization, Data curation, Investigation, Writing – review & editing, Supervision), and Philipp H. Schiffer (Conceptualization, Writing – review & editing, Supervision, Project administration, Investigation, Funding acquisition).

## SUPPLEMENTARY DATA

**Figure S1:** Comparison of body length of species.

**Figure S2:** Comparison of body length to greatest body diameter (a) of species.

**Figure S3:** Comparison of body length to neck length (b) of species.

**Figure S4:** Comparison of body length to tail length (c) of species.

**Figure S5:** Comparison of tail length to tail diameter (c’) of species.

**Figure S6:** Comparison of the distance of the vulva from the anterior of species.

**Figure S7:** Comparison of lip region width of species.

**Figure S8:** Comparison of stoma length of species.

**Figure S9:** Comparison of corpus length of species.

**Figure S10:** Comparison of isthmus length of species.

**Figure S11:** Comparison of basal bulb size of species.

**Figure S12:** Comparison of pharynx length of species.

**Figure S13:** Comparison of neck length of species.

**Figure S14:** Comparison of excretory pore to anterior end of species.

**Figure S15:** Comparison of body width at mid-body of species.

**Figure S16:** Comparison of body width at anus of species.

**Figure S17:** Comparison of tail length of species.

**Figure S18:** Comparison of the distance of the vulva to the anterior end of species.

**Figure S19:** Comparison of spicule length of species.

**Figure S20:** Comparison of gubernaculum length of species.

**Figure S21:** Blobplot of genome assembly of *P. einhardi* sp. nov.

**Figure S22:** Blobplot of genome assembly of *P. shuimeiren* sp. nov.

**Figure S23:** Blobplot of genome assembly of *P. nebliphilus* sp. nov.

**Figure S24:** K-mer spectra comparison generated with the K-mer Analysis Toolkit (KAT) comparing k-mers derived from PacBio HiFi sequencing reads to those present in the final genome assembly of *P. einhardi* sp. nov. (k = 27).

**Figure S25:** K-mer spectra comparison generated with the K-mer Analysis Toolkit (KAT) comparing k-mers derived from Oxford Nanopore sequencing reads to those present in the final genome assembly of *P. shuimeiren* sp. nov. (k = 27).

**Figure S26:** K-mer spectra comparison generated with the K-mer Analysis Toolkit (KAT) comparing k-mers derived from Oxford Nanopore sequencing reads to those present in the final genome assembly of *P. nebliphilus* sp. nov. (k = 27).

**Figure S27:** Circos plot visualizing pairwise alignments between the assemblies, displaying all scaffolds.

**Figure S28:** UpSet plot of orthologous gene clusters across species created with OrthoVenn3.

**Table S1**: Accession numbers of species downloaded from ENA for phylogenetic reconstruction.

## DATA AVAILABILITY

Sequencing reads will be deposited in the European Nucleotide Archive (ENA) under project number PRJEB75733. Scripts can be found on GitHub (https://github.com/lpettrich/PeinhardiAndTwoSisters), and the corresponding data are available on Zenodo (https://doi.org/10.5281/zenodo.17608368).

