## Supplementary Results for "*Panagrolaimus einhardi* sp. nov. and two sisters of fortune"

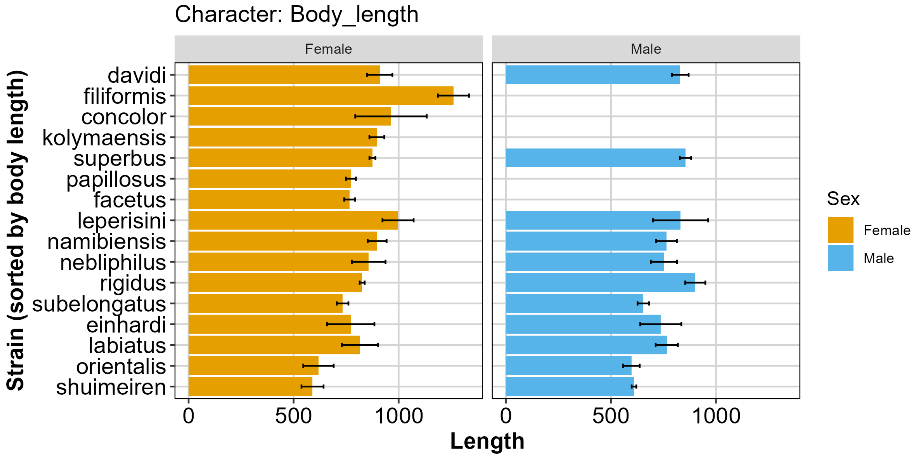


Figure S1: Comparison of body length (µm) of species.

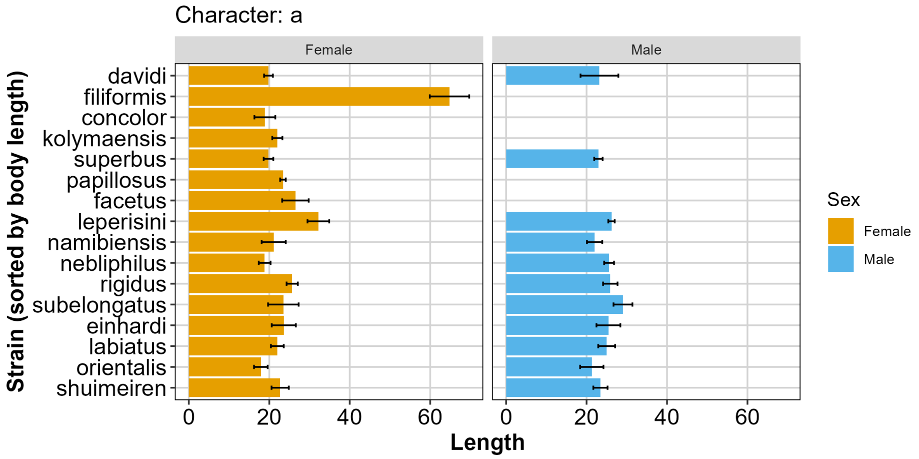


Figure S2: Comparison of body length to greatest body diameter (a) of species.

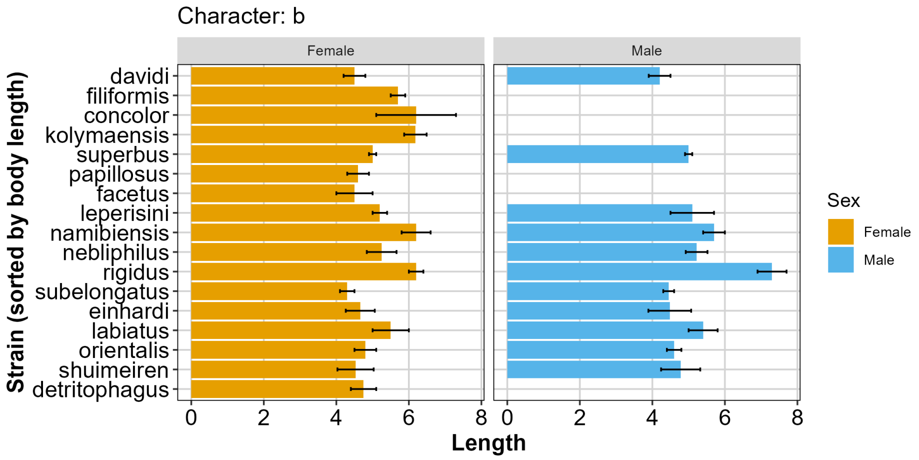


Figure S3: Comparison of body length to neck length (b) of species.

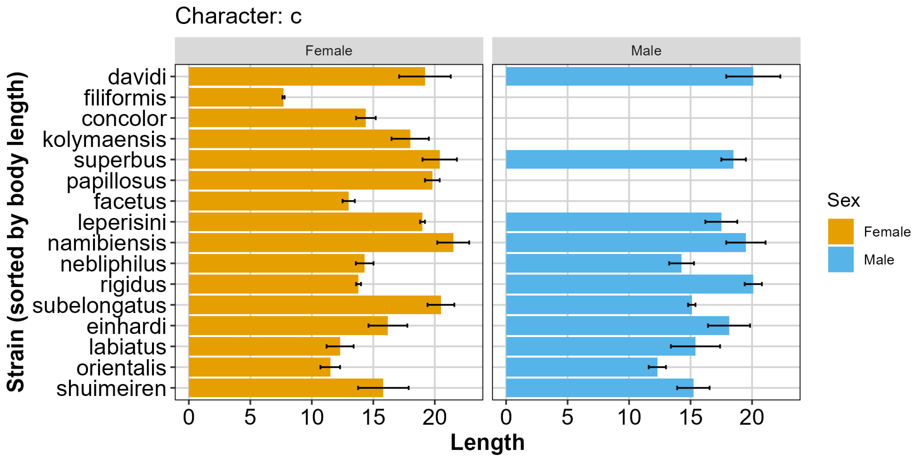


Figure S4: Comparison of body length to tail length (c) of species.

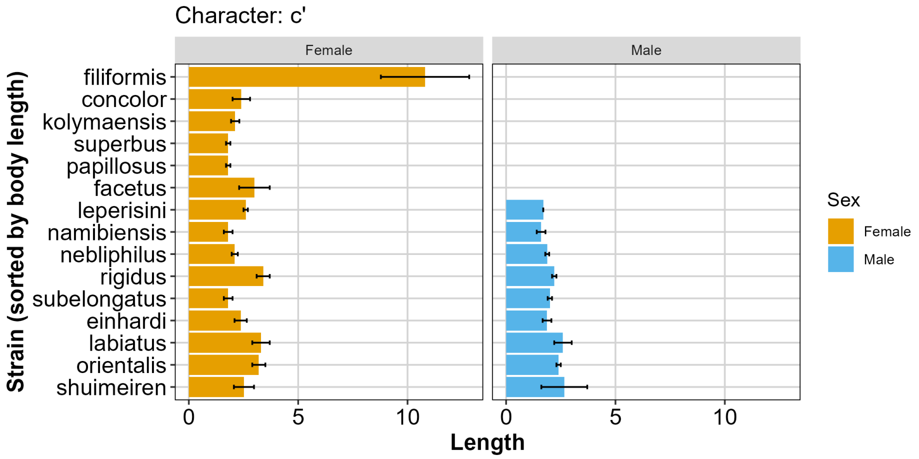


Figure S5: Comparison of tail length to tail diameter (c') of species.

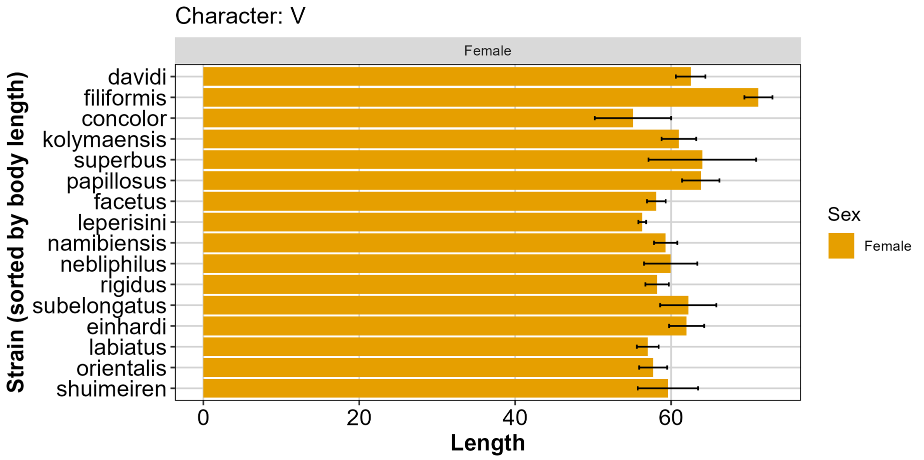


Figure S6: Comparison of the distance of the vulva from the anterior (%) of species.

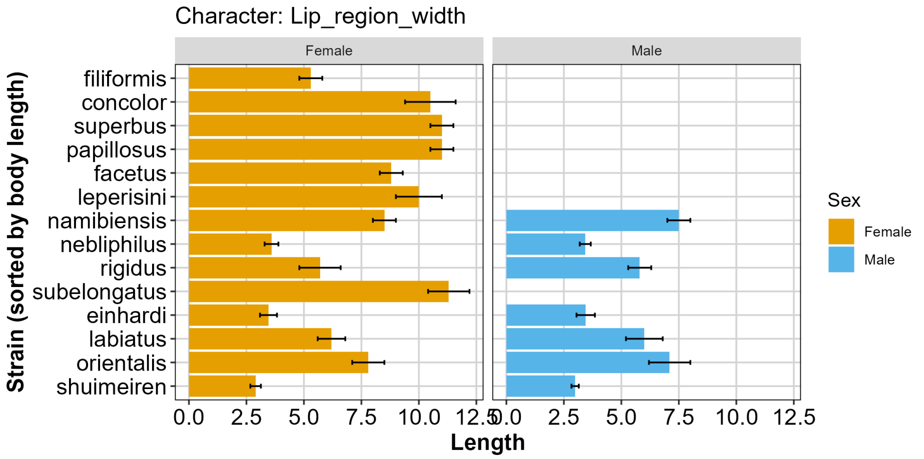


Figure S7: Comparison of lip region width (µm) of species.

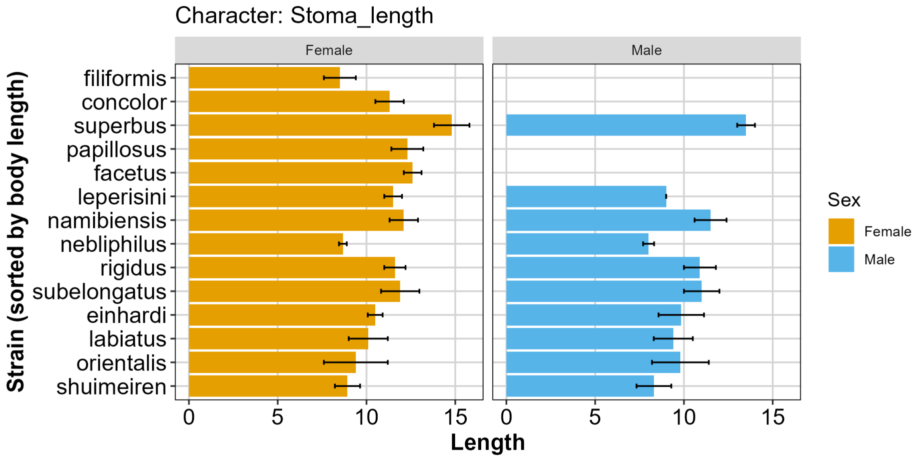


Figure S8: Comparison of stoma length (µm) of species.

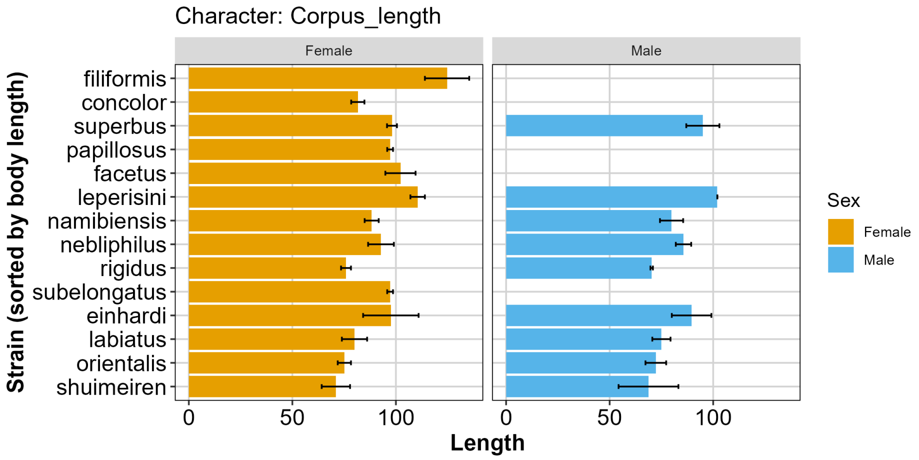


Figure S9: Comparison of corpus length (μm) of species.

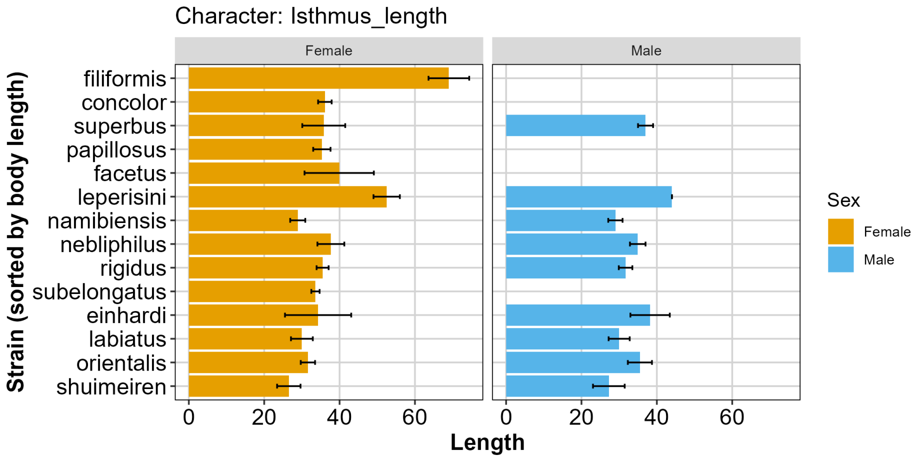


Figure S10: Comparison of isthmus length (μm) of species.

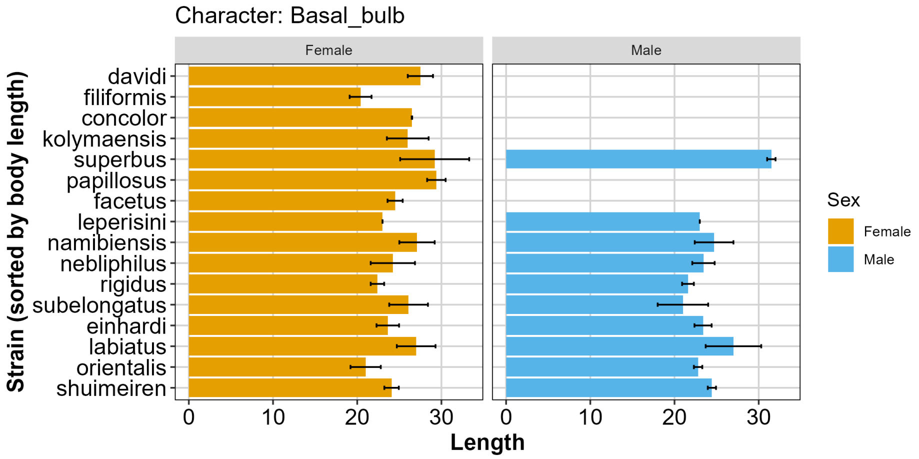


Figure S11: Comparison of basal bulb size (μm) of species.

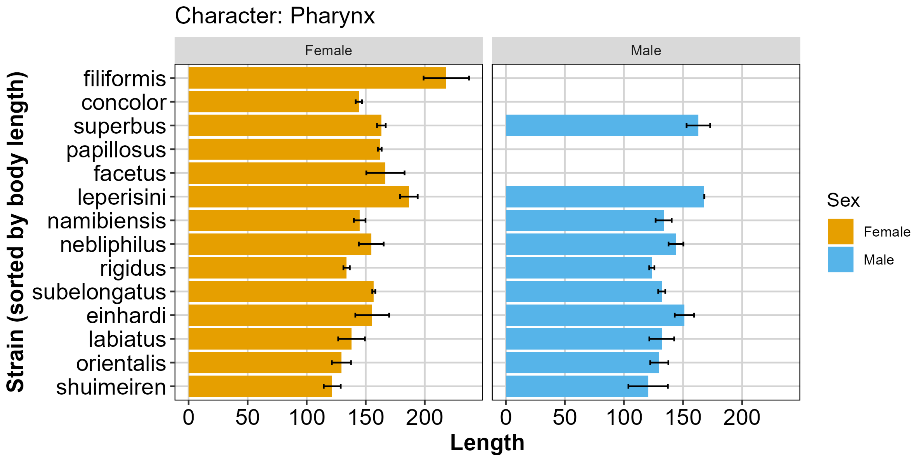


Figure S12: Comparison of pharynx length (µm) of species.

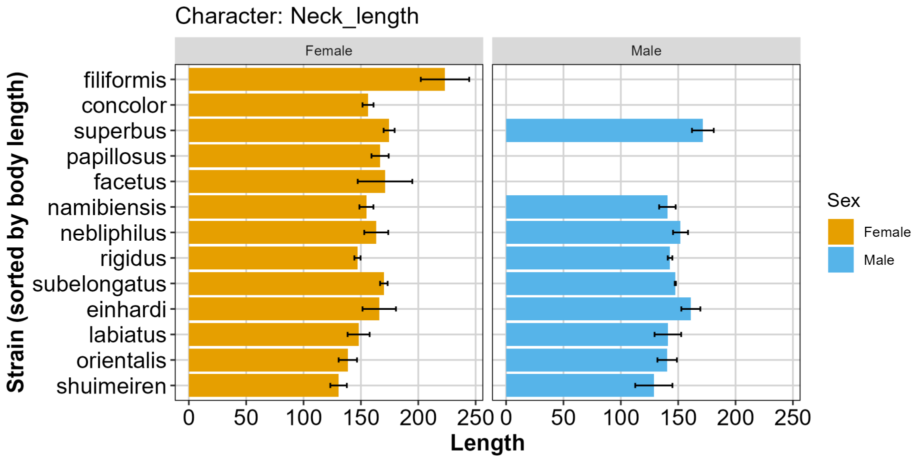


Figure S13: Comparison of neck length (µm) of species.

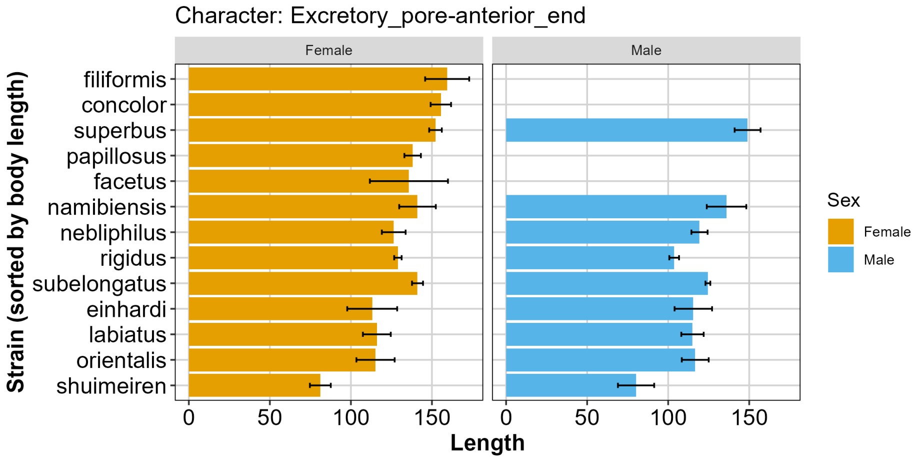


Figure S14: Comparison of excretory pore to anterior end (µm) of species.

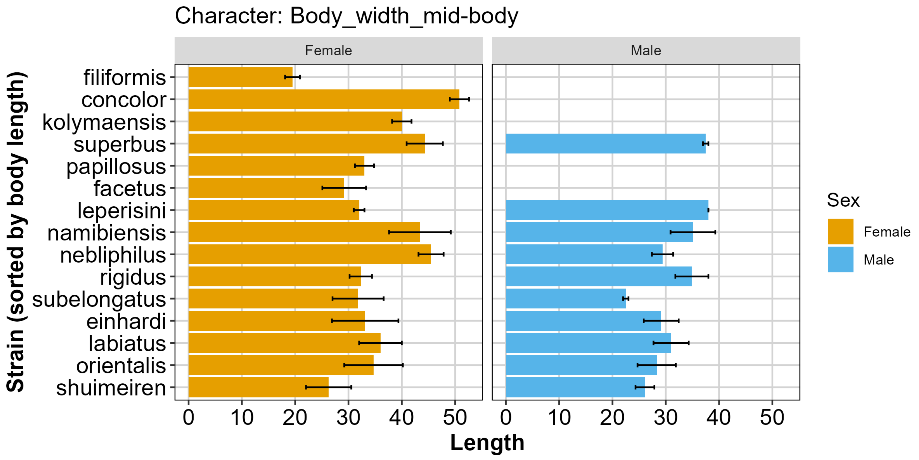


Figure S15: Comparison of body width at mid-body (µm) of species.

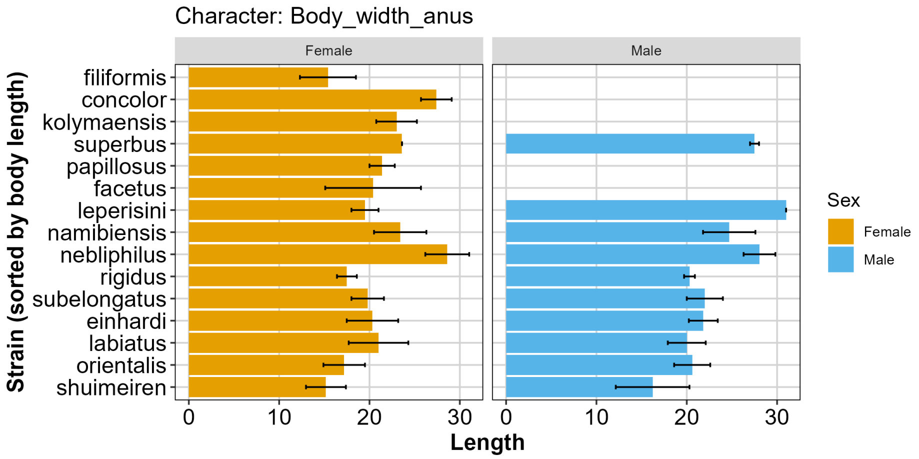


Figure S16: Comparison of body width at anus (µm) of species.

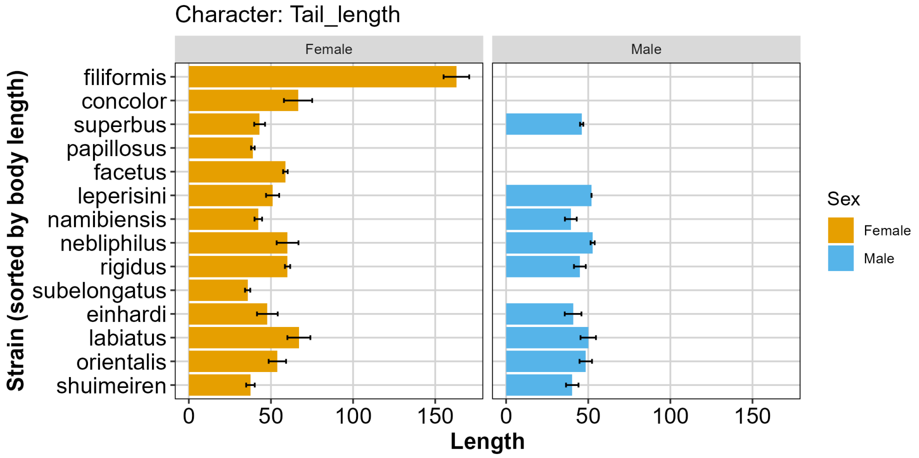


Figure S17: Comparison of tail length (µm) of species.

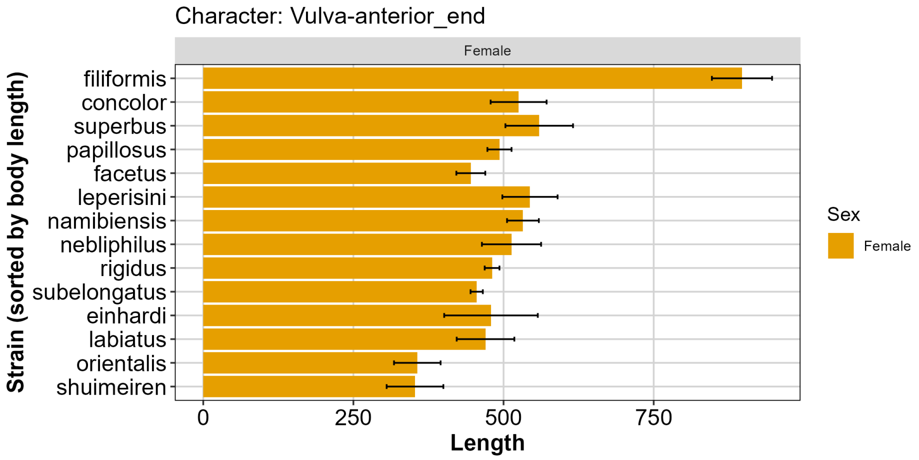


Figure S18: Comparison of the distance of the vulva to the anterior end (µm) of species.

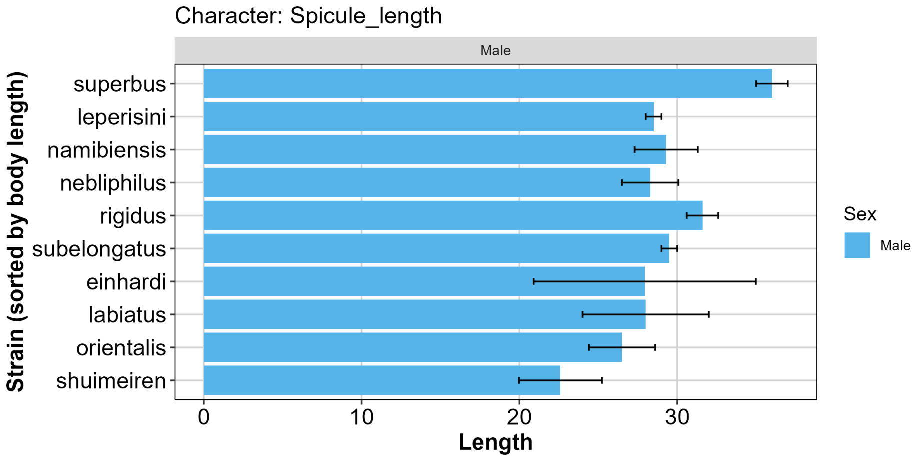


Figure S19: Comparison of spicule length (µm) of species.

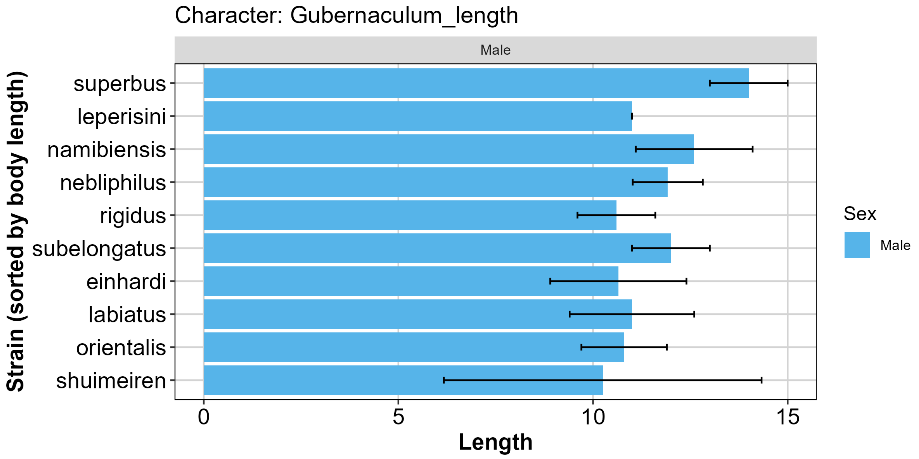


Figure S20: Comparison of gubernaculum length (µm) of species.

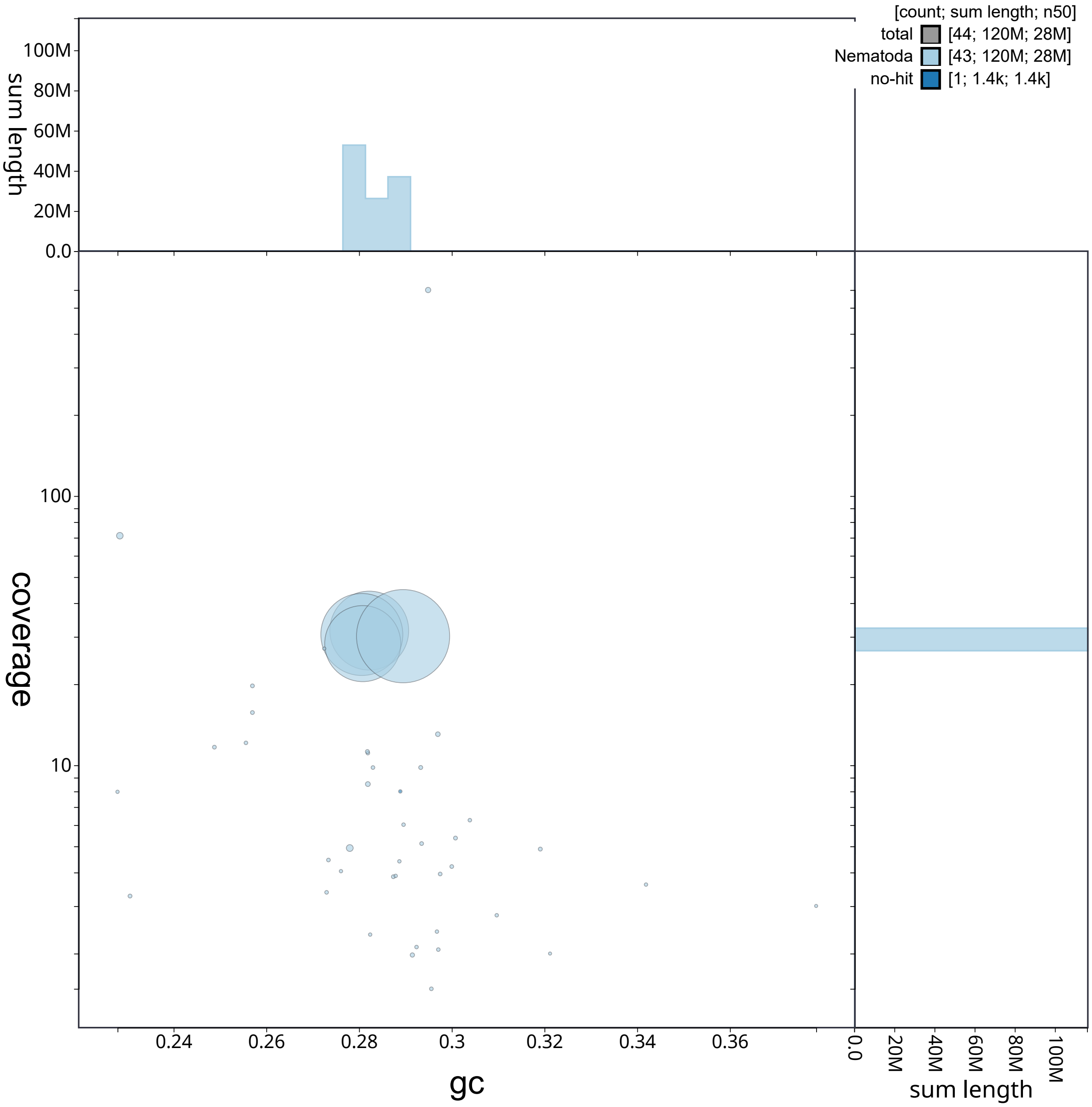


Figure S21: Blobplot (Challis et al., 2020) showing GC content versus long-read sequencing coverage (PacBio HiFi) for scaffolds in the genome assembly of *P. einhardi* sp. nov. Scaffolds are colored according to taxonomic assignment and the size refers to read length.

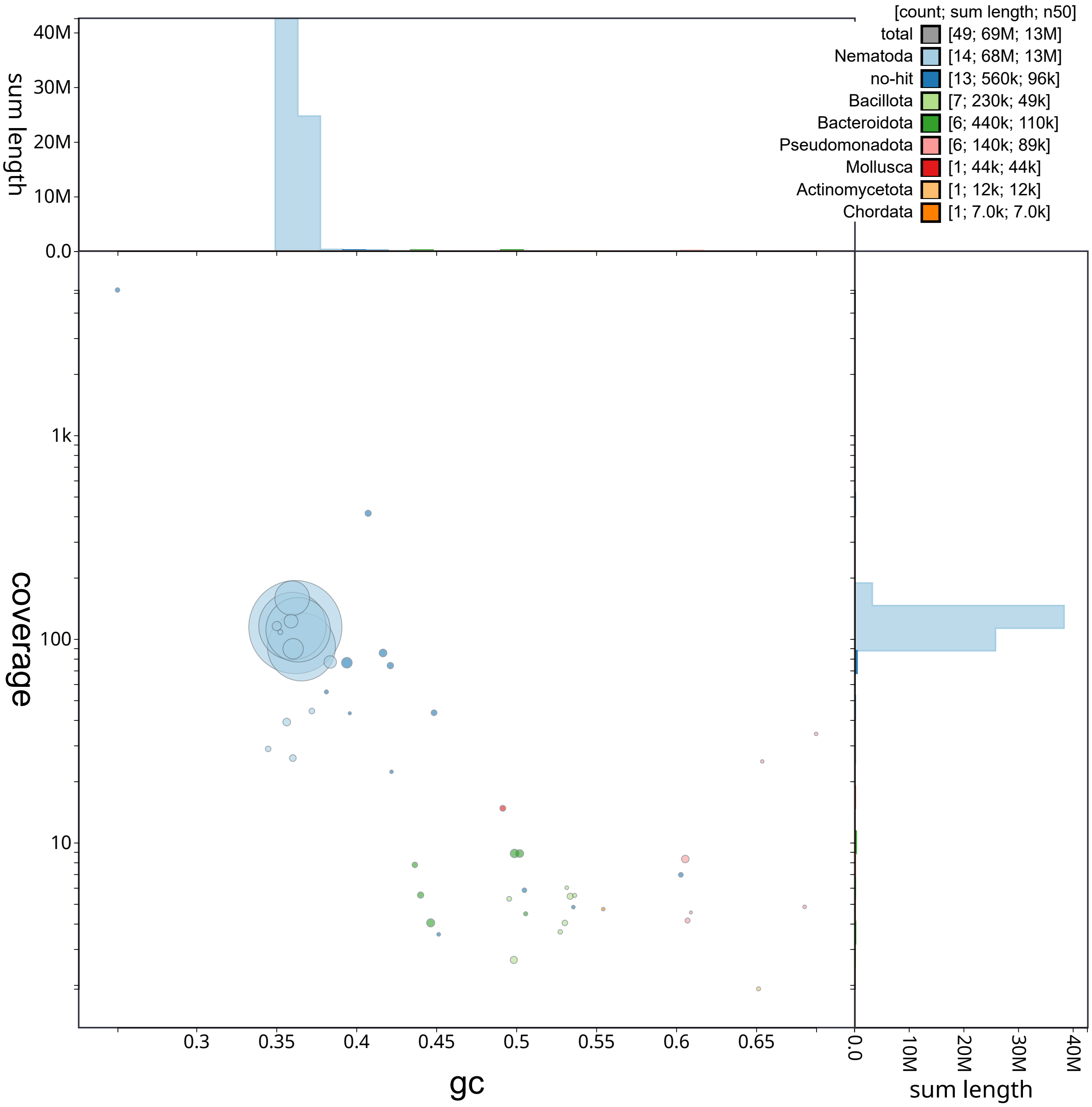


Figure S22: Blobplot (Challis et al., 2020) showing GC content versus long-read sequencing coverage (Oxford Nanopore) for scaffolds in the genome assembly of *P. shuimeiren* sp. nov. Scaffolds are colored according to taxonomic assignment and the size refers to read length.

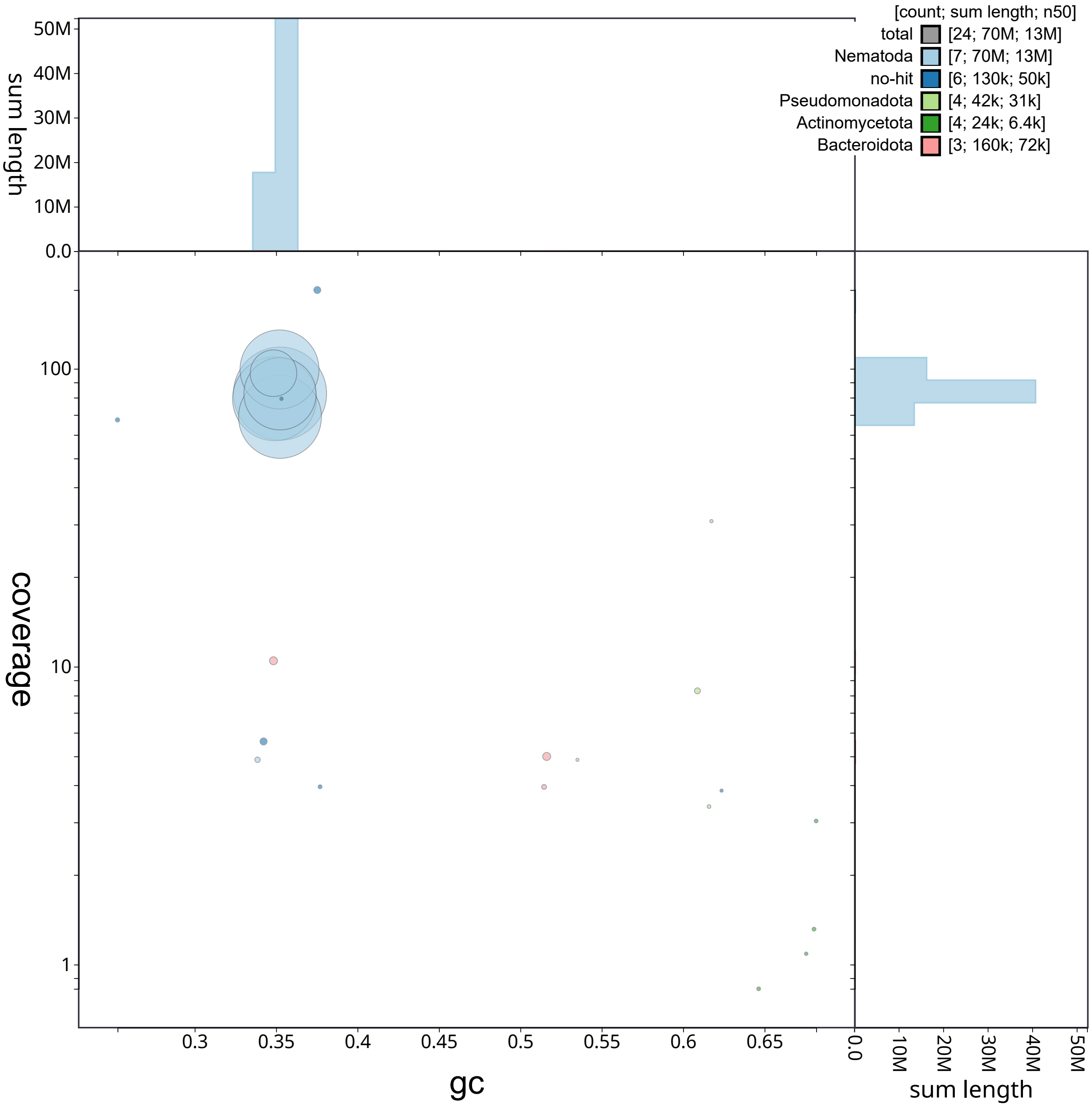


Figure S23: Blobplot (Challis et al., 2020) showing GC content versus long-read sequencing coverage (Oxford Nanopore) for scaffolds in the genome assembly of *P. nebliphilus* sp. nov. Scaffolds are colored according to taxonomic assignment and the size refers to read length.

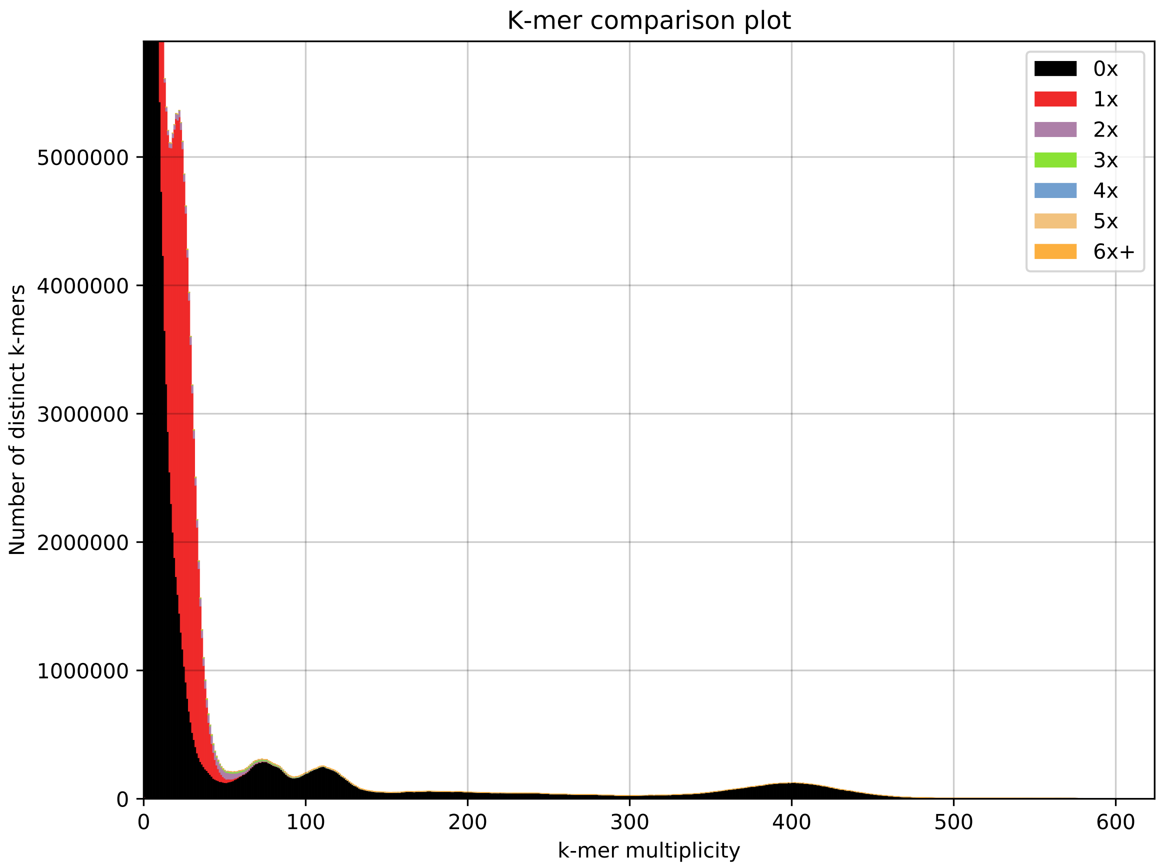


Figure S24: K-mer spectra comparison generated with the K-mer Analysis Toolkit (KAT) (Mapleson et al., 2016), comparing k-mers derived from PacBio HiFi sequencing reads to those present in the final genome assembly of *P. einhardi* sp. nov. (k = 27).

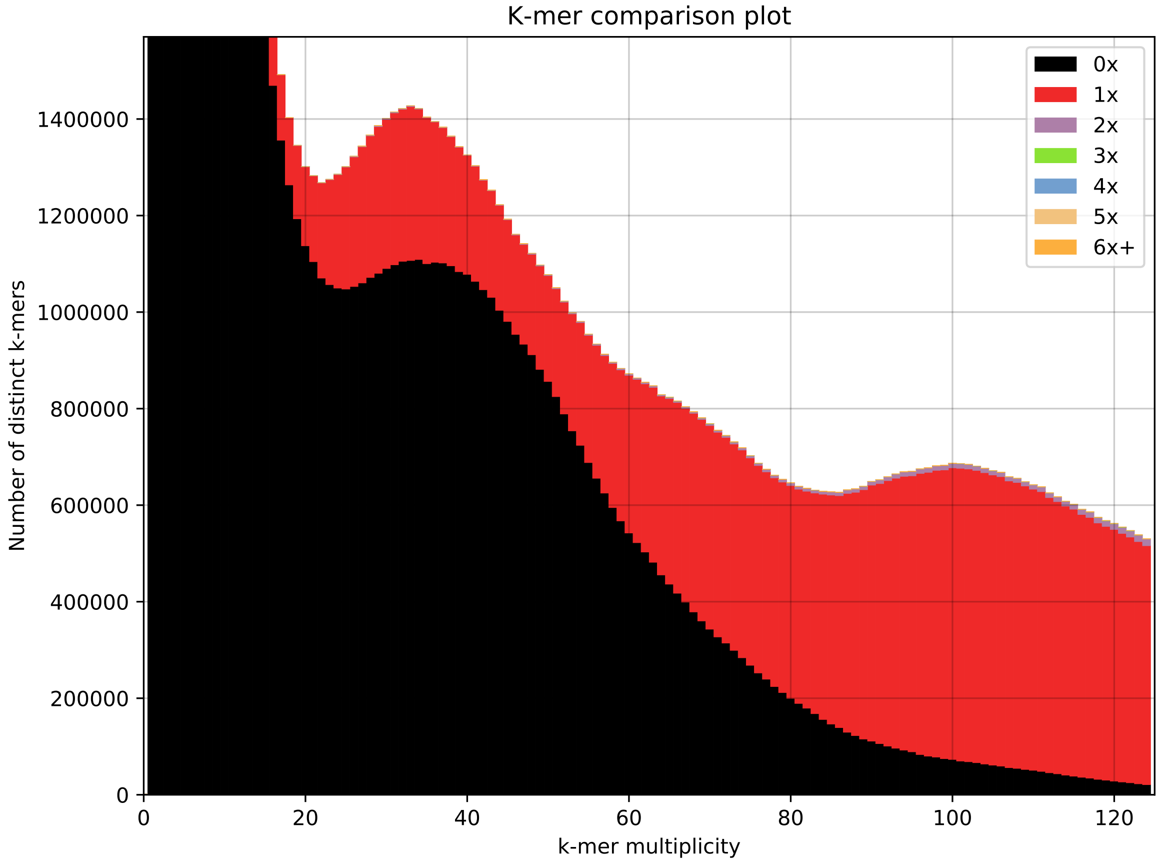


Figure S25: K-mer spectra comparison generated with the K-mer Analysis Toolkit (KAT) (Mapleson et al., 2016), comparing k-mers derived from Oxford Nanopore sequencing reads to those present in the final genome assembly of *P. shuimeiren* sp. nov. (k = 27).

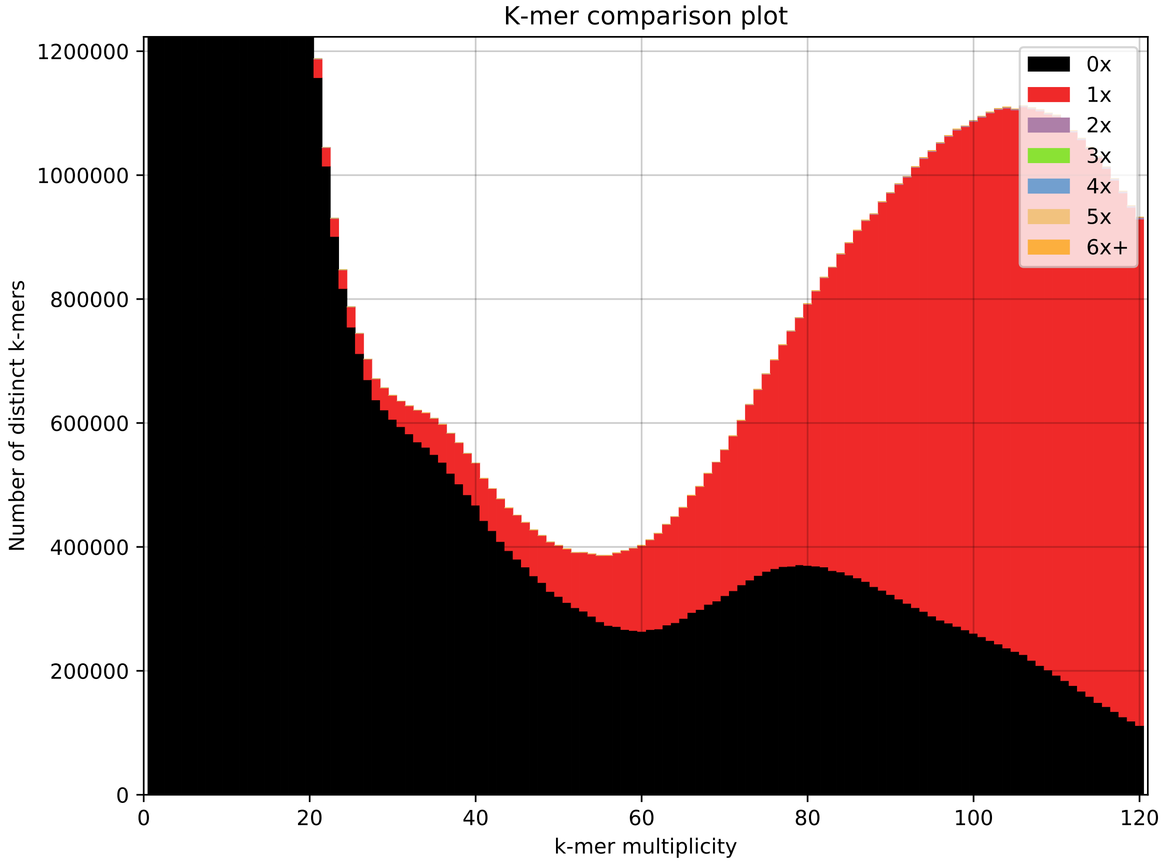


Figure S26: K-mer spectra comparison generated with the K-mer Analysis Toolkit (KAT) (Mapleson et al., 2016), comparing k-mers derived from Oxford Nanopore sequencing reads to those present in the final genome assembly of *P. nebliphilus* sp. nov. (k = 27).

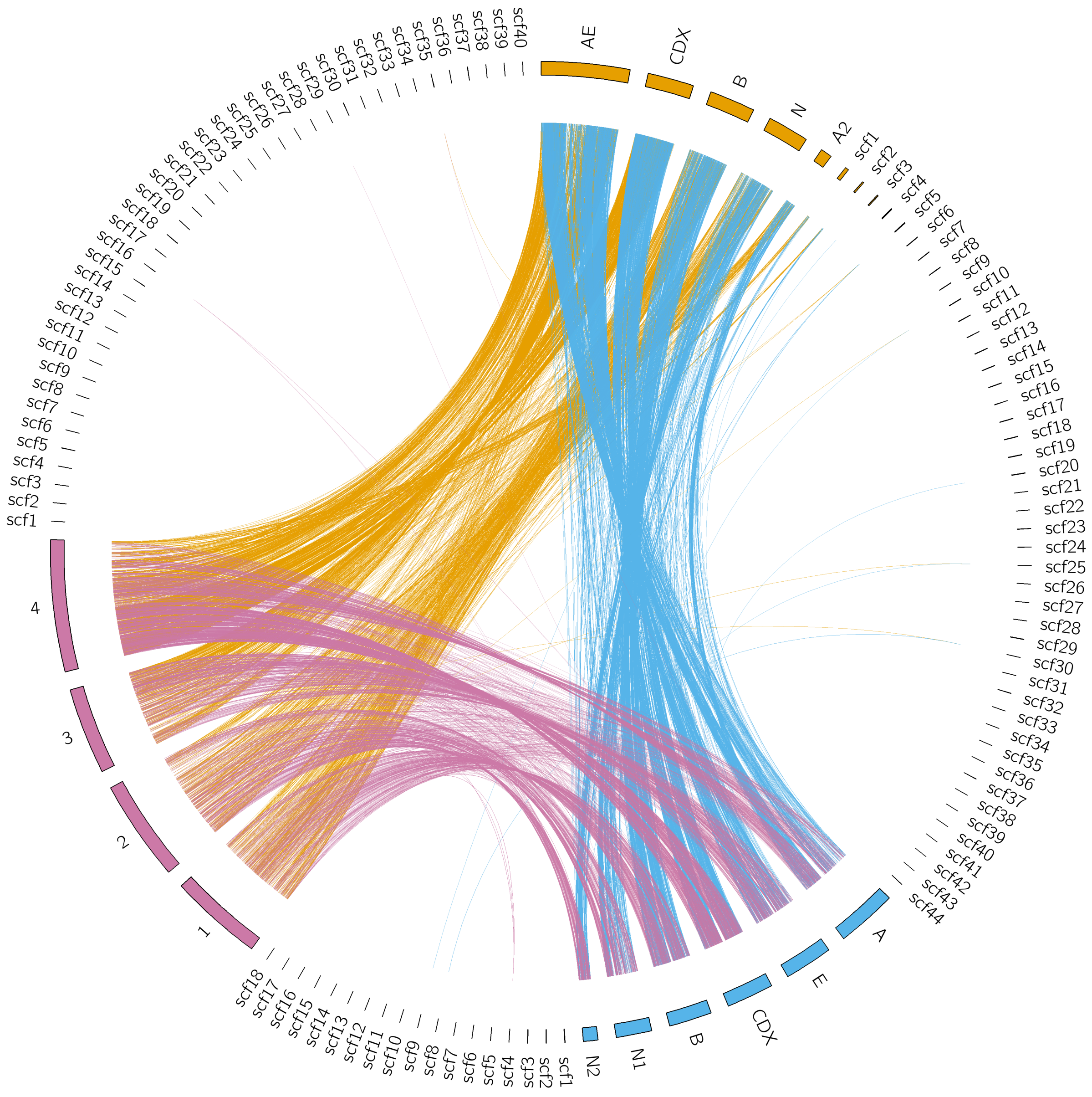


Figure S27: Circos plot (Krzywinski et al., 2009) visualizing pairwise alignments between the assemblies displaying all scaffolds. The colors indicate the species: Red = *P. einhardi* sp. nov., orange = *P. shuimeiren* sp. nov., blue = *P. nebliphilus* sp. nov.

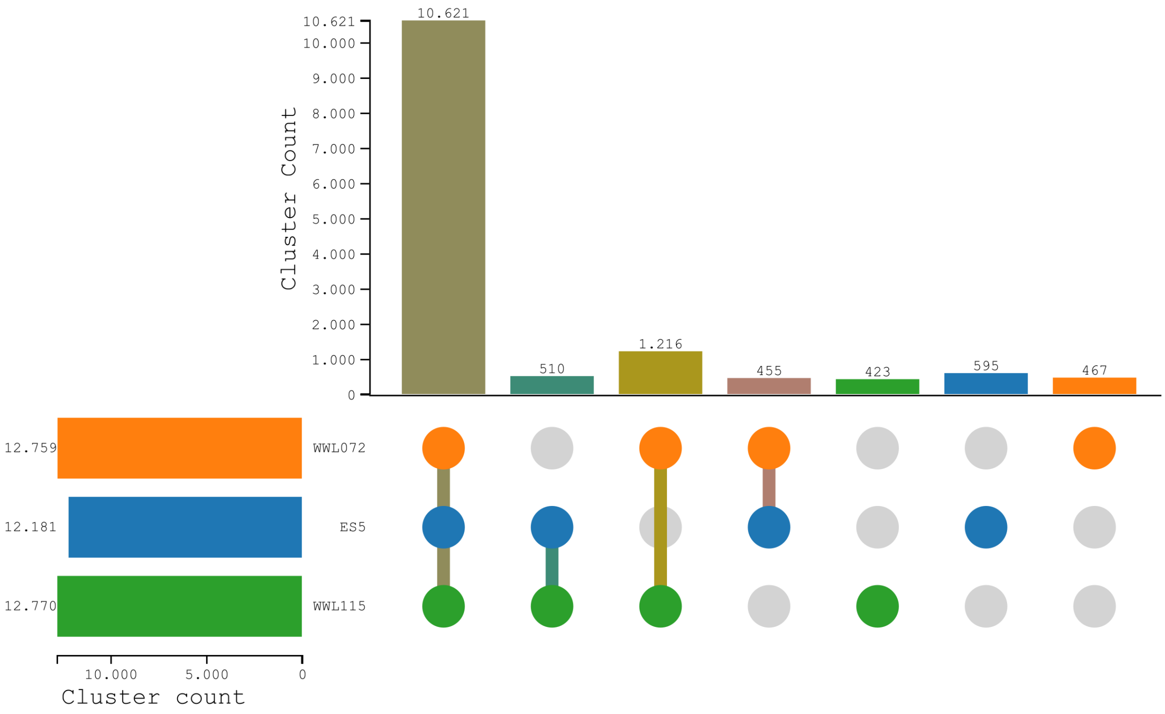


Figure S28: UpSet plot of orthologous gene clusters across species created with OrthoVenn3 (Sun et al., 2023). Total cluster count for each species is given (WWL072 = *Panagrolaimus nebliphilus* sp. nov; ES5 = *Panagrolaimus einhardi* sp. nov.; WWL115 = *Panagrolaimus shuimeiren* sp. nov.) in horizontal bars on the left. Vertical bars indicate the number of clusters in each intersection, where connected dots below specify which species are included.

Table S1: Accession numbers of species downloaded from ENA for phylogenetic reconstruction using ultra-concserved elements (UCEs).

| **Name** | **Accession** |
|---|---|
| *Acrobeloides maximus* | GCA_964212105 |
| *Acrobeloides obliquus* | GCA_034698545 |
| *Acrobeloides thornei* | GCA_034699885 |
| *Halicephalobous* sp. NKZ332 | GCA_009761265 |
| *Halicephalobus mephisto* | GCA_009193035 |
| *Panagrellus redivivus* | GCA_000341325 |
| *Panagrolaimus davidi* | GCA_901779475 |
| *Neocephalobus halophilus* BSS8 | GCA_964187885 |
| *Panagrolaimus* sp. LJ2400 | GCA_024447215 |
| *Panagrolaimus* sp. LJ2406 | GCA_024447205 |
| *Panagrolaimus* sp. LJ2414 | GCA_024447195 |
| *Panagrolaimus* sp. PS1159 | GCA_963922195 |
| *Panagrolaimus* sp. PS1579 | GCA_901779485 |
| *Panagrolaimus superbus* | GCA_901766145 |
| *Propanagrolaimus* sp. JU765 | GCA_901765185 |
